# Multi-scale Modeling and Experimental Data Enable Structural Models of the *Escherichia Coli* Peptidoglycan to be Constructed on the Whole-Cell Scale

**DOI:** 10.1101/2023.06.04.543634

**Authors:** Zachary J. Wehrspan, Adrian H. Elcock

## Abstract

The peptidoglycan (PG) layer of *Escherichia coli* is a single, interconnected gigaDalton molecule that is the largest in the cell. Experimental studies have established a number of the PG’s properties, and previous computational studies have simulated aspects of its behavior on sub-cellular scales, but none have fully modeled the PG’s compositional heterogeneity and no models have yet been constructed on the whole-cell scale. Here we use a combination of computational modeling approaches to construct whole-cell PG models at a resolution of one coarse-grained (CG) bead per glycan that are consistent with a wide variety of available experimental data. In particular, we derive plausible glycan strand length distributions for the polar and cylindrical regions of the cell that cover the full range of possible strand lengths and that are consistent with all available experimental data. In addition, we develop stochastic simulation code that explicitly models a cross-linking experiment from the literature that has a direct bearing on the extent to which Braun’s lipoprotein (Lpp) is partitioned between periplasmic and surface-exposed locations. We then use all of these data as inputs to a new computer code, *PG_maker*, which builds CG models of the PG on a whole-cell scale in under an hour. Finally, we use the resulting 3D models as a basis for: (a) estimating pore size distributions – which, despite the idealized nature of the models, are shown to be in surprisingly good agreement with experimental estimates – and (b) calculating the effects of the large numbers of periplasmic Lpps on the ability of freely diffusing proteins to access the compartment that lies between the PG and the outer membrane. The ability to combine a wide range of experimental data into structural models that are physically realizable in 3D helps to set the stage for performing simulations of the PG on the whole-cell scale in the near future.

## Introduction

The peptidoglycan (PG) layer of Gram-negative bacteria has long been known to be critical for maintain their structural integrity (1). The PG is a meshwork of glycan strands (with a wide distribution of lengths) that, to a first approximation, can be thought of as being roughly arranged in hoops that wrap around the long axis of the cell and that are cross-linked, to varying degrees, by the enzyme-catalyzed formation of covalent bonds between peptide sidechains that protrude from the glycans. Importantly, cellular PG is also covalently attached to thousands of copies of Braun’s lipoprotein (Lpp), a trimeric protein whose N-termini are tripalmitoylated and resident in the outer membrane (OM) (2,3). Lpp is the only protein known to be covalently connected to the PG (4), and this connection has been shown to be a key determinant of the cell’s mechanical properties (5). In addition, however, a number of other proteins are known to bind the PG non-covalently. These include two very abundant proteins – the outer-membrane porin OmpA (6), and the lipoprotein Pal (7), an integral component of the Tol-Pal complex that plays an important role in the complicated choreography that accompanies cell division (8) – as well as a variety of SPOR domain-containing proteins that bind to peptide-denuded PG at the site of cell division (9).

Over the years, an impressive amount of information on the chemical composition of the PG has been obtained by experimental methods. Glycan strands are known to consist of alternating sequences of the monosaccharides *N*-acetylglucosamine (GlcNAc) and *N*-acetylmuramic acid (MurNAc), with the final glycan in each strand being a 1,6-anhydro MurNAc. The lengths of the glycan strands can vary, however, and several efforts have been made to determine the distribution (e.g. (10–12)): average strand lengths of ∼25 disaccharide (DS) units have been reported, and at a more detailed level, the relative populations of all strand lengths lower than 30 disaccharide (DS) units have been largely resolved. Other studies have focused on the chemical composition of the PG’s peptide components, with relative populations of the various possible peptide sidechains and inter-sidechain cross-links quantified (11,13,14). Some of these experimental studies have identified potentially significant differences between the chemical composition of the PG laid down in the cylindrical and the polar regions of the cell (11). Finally, experimental studies have sought to determine in detail the nature of the relationship between the PG and Lpp, with a particularly provocative study suggesting that a substantial fraction of Lpp might not, in fact, be covalently connected to the PG but might instead be exposed on the cell surface (15).

In addition to these insights into the chemical composition of the PG, experimental techniques have also highlighted physical properties and features of the PG. AFM imaging of patches of PG sacculi isolated from cells, for example, has been used to determine distributions of pore sizes (16,17) and to show that individual glycan strands can be as long as 200 disaccharide units. Very recently, AFM imaging has also been used in an attempt to identify Lpp molecules covalently attached to PG sacculi (18); the size and numbers of Lpp “lumps” observed in those experiments have been tentatively suggested as being representative of clusters of Lpp molecules. A quite different experimental technique that has been used to image PG *in situ* is cryoelectron tomography, which has been shown to be capable of identifying glycan strands, and has provided estimates of both the thickness of the PG layer (it appears to be mostly single-layered, for example (19)) and the average distance between the PG and both the outer and inner membranes (20).

Additional insights into aspects of PG behavior have accrued from the use of molecular simulation strategies. At one end of the spectrum have been atomically detailed molecular dynamics simulations; these have included pioneering studies by the Gumbart group simulating the mechanical properties of the PG at the atomic level (21,22) and a series of interesting studies from the Khalid group that have provided intriguing insights into the ways in which PG might interact with periplasmic and outer membrane proteins ((23–25); see also (20)). At the other end of the spectrum, a number of innovative coarse-grained (CG) simulation models have been developed for use in large-scale simulations by the Wingreen and Huang groups (26–28) and by the Jensen group (29,30). These models have been used to study the PG’s role in maintaining cell shape and to propose and test mechanisms by which the PG might be synthesized and remodeled during cell growth and division; they operate at a resolution of one coarse-grained bead for every two or four glycans and have been used to reach length-scales that approach perhaps 15 % of the length of a typical bacterial cell. Finally, the Ayappa group has developed an intermediate-resolution CG model of the PG (31), suitable for use with the popular MARTINI force field (32–35), that offers considerable promise for simulating the PG on large, but probably still sub-cellular scales.

One major question that arises from all of the above studies is the extent to which the information derived from them can be combined into a single, self-consistent model that provides a structural view of the PG on a whole-cell scale. We aim to achieve this here by exploiting a variety of simulation and modeling strategies, each subject to constraints imposed by corresponding experimental data, and each focused on the PG of *Escherichia coli*. These efforts focus on filling in those aspects of the puzzle that have so far remained unresolved experimentally, and finding ways to combine the various sources of data in ways that ensure they remain mutually consistent. The insights obtained from individual stages are then synthesized and used as input to new computer code that rapidly builds CG molecular models of the PG.

The resulting models have a level of structural resolution that is intermediate between the highly detailed views obtained from previous atomistic simulations and the less detailed views obtained from previous large-scale CG models, but that is implemented on a whole-cell scale that is, to our knowledge, unprecedented. Importantly, when used to build an idealized model of the PG of *E. coli* cells growing in rich defined media, the code directly predicts a total copy number for Lpp that matches well with a ribosome-profiling-based experimental estimate (36) that was not used during model construction.

To demonstrate the potential power of having whole-cell PG models, we analyze two aspects of the models that are likely to have ramifications for *E. coli*’s physiology. First, we use image processing software to determine pore size distributions in the whole-cell model, finding good agreement with the Foster group’s finding that the frequency with which pores of a given size are present decays approximately exponentially with increasing pore area. Second, we use CG models of representative proteins to measure the effects of the highly crowded Lpp connections on the ease with which periplasmic proteins can access the periplasmic space that lies between the PG and the OM. We find that the density of Lpp is sufficiently high in the whole-cell model that the largest proteins are thermodynamically disfavored from accessing the OM-proximal compartment even if they can find a way through the PG mesh. The availability of a method for rapidly building whole-cell PG models sets the stage for future work aimed at modeling the conformational dynamics of the PG layer on the cellular scale.

## Results

While much is already known experimentally about the chemical composition of the PG (see Introduction), there are some critical issues for which information is either scarce or controversial. Two key areas where important questions remain without definitive answers include: (1) the complete distribution of the lengths of glycan strands within the PG, and (2) the relative populations of Lpp that are: (a) resident in the periplasm, and that are therefore available for covalent attachment to the PG, or (b) exposed on the cell surface. To obtain answers to these questions we make use of a combination of statistical and simulation approaches; we then integrate these results into computer code that builds coarse-grained models of the PG on the whole-cell scale in under an hour.

### Modeling the distribution of glycan strand lengths

The relative populations of glycan strand lengths ranging from 1 to 30 DS units have been quantified using high performance liquid chromatography (HPLC) by Obermann and Höltje (11); these data have already been used previously as a basis for generating large-scale PG models by the Wingreen group (26). Obermann and Höltje’s data are for a strain of *E. coli* that exhibits both normal and “minicell” phenotypes; the latter was used by those authors as a proxy for understanding the properties of PG in the polar regions of the cell, and subtracting a suitably weighted version of its glycan length distribution from that of wild-type cells makes it possible to estimate the glycan length distribution in the cylindrical regions of the cell (see Materials & Methods). While the distribution of glycan strand lengths is known experimentally for strands of length ≤ 30 DS units, the distribution for longer strands is not known in detail and our first goal, therefore, is to develop a model that allows us to estimate it. To do this we make use of the following additional experimental information. First, while the work of Obermann and Höltje data does not provide a detailed distribution beyond 30 DS units, it does tell us that the combined population of DS units from all strands longer than 30 DS units is 13%. Second, the same study also reports an estimate of the *mean* length of a glycan strand: 27.8 DS units. Obermann and Höltje calculated the latter number by exploiting the fact that each glycan strand terminates in a 1,6-anhydro MurNAc residue and dividing the total measured glycan mass by the measured mass of 1,6-anhydro MurNAc. Importantly, therefore, this estimate was obtained using a method that was independent of the way the glycan length distribution was measured. Estimates of the mean glycan strand length in *E. coli* made by other groups also typically amount to ∼30 DS units (13,14,37–39). Third, as noted in the Introduction, we know from atomic force microscopy (AFM) experiments performed by the Foster group that individual glycan strands can be very long: strands have been identified by AFM that appear to be at least as long as 200 DS units (17), which if fully extended would cover approximately 1/25 of the circumference of a typical *E. coli* cell. The existence of very long strand lengths is consistent with the highly processive nature of the elongasome machinery that is primarily responsible for building them: single-molecule tracking studies of MreB filaments performed by the Garner group (40), for example, have indicated that multiple complete circuits of the cell circumference can be performed during the lifetime of a single filament.

In support of the AFM experimental data, the following simple calculation indicates that the unresolved population in Obermann and Höltje’s experiments must consist of a significant number of very long strands. The mean length of the combined population of strand lengths ≤ 30 DS units reported by Obermann and Höltje is 8.3 DS units (11). We know that this short-strand population comprises 87% of the total population, so the remaining 13% of the population (see above) must be of a sufficient length to bring the mean strand length up to 27.8 DS units. In other words, if the “missing” long-strand population has a mean length of α DS units, then (0.87 × 8.3) + (0.13 × α) = 27.8. Solving for α, we obtain the result α = 158 DS units. We should not be surprised, then, if the “true” underlying distribution of glycan strand lengths contains a substantial population of long glycan strands.

Given these lines of evidence, we have attempted to derive a glycan length distribution that simultaneously matches: (a) Obermann and Höltje’s experimental distribution for strand lengths ≤ 30 DS units, (b) their reported relative populations of all strands with ≤ 30 DS units and all strands with > 30 DS units, and (c) their reported mean strand lengths. Since we found that these criteria could not be simultaneously satisfied by assuming that the distribution follows a simple monotonic decay, we assumed instead that the strand length distribution is bimodal, and used the experimentally reported values for all strands with ≤ 30 DS units (supplemented with a power-law fit of those data to predict the distribution for strands up to 37 DS units) together with a Gaussian distribution of fixed width centered at a much longer strand length (see Materials & Methods). With the width of the Gaussian set arbitrarily at 30 DS units, optimizing the fit to all three criteria gives Gaussians centered at 165 and 260 DS units for the cylindrical and polar regions of the cell, respectively; the positions of these Gaussian centers vary only very modestly when the widths of the Gaussians are instead set to 15 and 60 DS units (see Figure S1). The final complete glycan length distributions that we obtain for the cylindrical and polar region of the cell are shown in Figure 1a; plots focusing on the short and long strand regions of the distributions are shown in Figures 1b and 1c, respectively. To the extent that minicells provide a good model of polar PG, the plots shown in Figure 1 suggest that polar PG has a greater proportion of very short glycan strands relative to cylindrical PG and a minor population of long glycan strands whose lengths are typically greater than those found in cylindrical PG.

**Figure 1.**
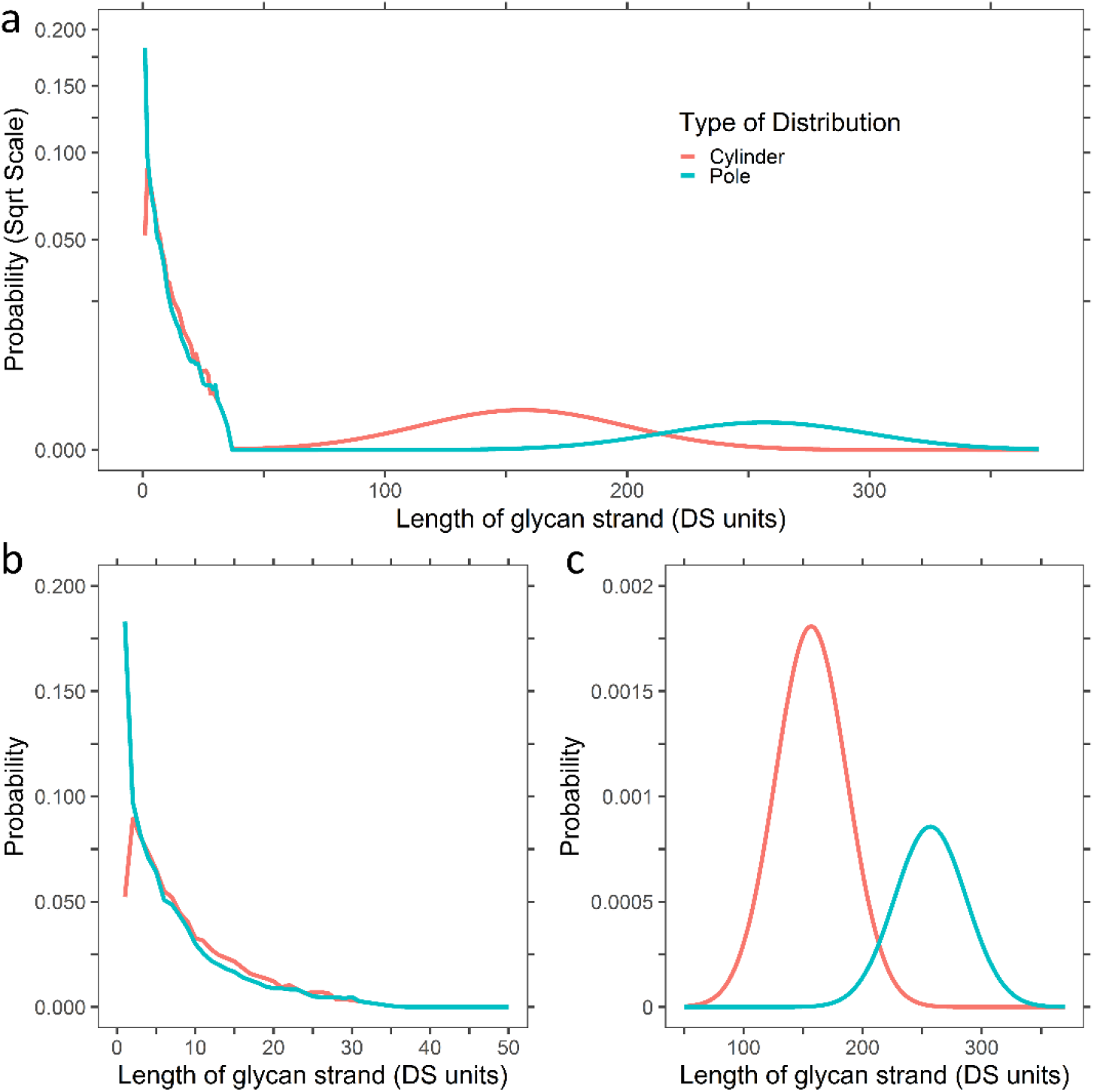
Derived glycan strand length distributions for the cylindrical (red) and polar regions of the cell (blue). (a) The full distribution for strands ranging from 1 to 370 DS units is plotted on a square root scale. (b) A zoomed in view of the distribution for 1 to 50 DS units. (c) A zoomed in view of the distribution for 50 to 370 DS units.

### Modeling the relative populations of Lpp in the periplasm and on the cell surface

A second area where a significant gap in our knowledge of PG structure exists concerns the intracellular distribution of Braun’s lipoprotein (Lpp). This highly abundant trimeric protein, one end of which resides in the OM via its tripalmitoylated N-terminal cysteine residues, is a critical component to include in any true whole-cell PG model since its C-terminal lysine residues can be covalently linked to the peptide sidechains of the PG (2). The conventional “textbook” view of Lpp places it exclusively in the periplasm (e.g. (41–43)) and directly connected to the peptide component of the PG, but intriguing data from the Silhavy group (15) has indicated that a substantial fraction of the Lpp population (as much as two-thirds of the total population) may instead be exposed on the cell surface. Since these two viewpoints, taken at face value, conjure up very different structural pictures of the PG and Lpp’s relationship to it, it seems important to determine whether the data underlying them can be reconciled to yield a unified model. To this end, we sought to develop a computer simulation approach capable of replicating the critical experimental data that led to the suggestion that the majority of the Lpp population is surface-exposed.

The Silhavy group’s study contains several lines of evidence that suggest that at least some fraction of Lpp is exposed on the cell surface, but the argument that the majority of the Lpp is located there is largely based on a chemical cross-linking experiment performed *in vivo*. Throughout that work, much of the discussion centers on differentiating between a so-called “free” form of Lpp (which comprises those Lpp chains that are not covalently connected to the PG) and a “bound” form of Lpp (which comprises those Lpp chains that are covalently connected to the PG). To support the idea that the free and bound forms of Lpp occupy different subcellular compartments, Cowles et al. compared the effects of the chemical cross-linking agent disuccinimdyl suberate (DSS) on cells expressing wild-type Lpp with its effects on cells expressing an Lpp mutant (ΔK_58_) mutant that is incapable of being covalently connected to the PG (and for which the population of “bound” Lpp must, therefore, be zero). The observation that similar levels of trimeric cross-linked Lpp were obtained with both cell lines was argued as being consistent with the notion that essentially all of the free form of the Lpp is surface-exposed (15).

We wondered whether the cross-linking data presented in Cowles et al. might be subject to alternative interpretations that might be more in line with the conventional view of Lpp’s subcellular location. We therefore developed stochastic simulation code that is intended to mimic the cross-linking experiment as closely as possible. Briefly summarized, the cross-linking experiment itself proceeds as follows: (1) cells are incubated with the cross-linker DSS for twenty minutes, (2) Lpp chains that are *not* covalently linked to the PG are isolated, and (3) they are separated into uncross-linked monomers, cross-linked dimers, and cross-linked trimers. Importantly, Lpp chains that are already covalently attached to the PG directly are assumed to not be quantified by the experiment, and nor are Lpp chains that, while not covalently attached to the PG directly, are themselves cross-linked by DSS to Lpp chains that are. In developing a simulation scheme that attempts to model the above experiment at a quantitative level, our first goal was to try to quantify the relative populations of monomers, dimers, and trimers from the immunoblots shown in Figure 6 of Cowles et al. (15); we did this using a straightforward analysis with ImageJ/Fiji (44,45) (see Materials & Methods). Plots showing the relative intensities of the immunoblots as a function of the distance traveled down the gel are shown in Figure S2a and S2b for wild-type Lpp and for the ΔK_58_ Lpp mutant, respectively. The latter contains a sufficiently high background signal for the oligomeric species that it seems impossible to quantify the relative populations with any degree of certainty; it also appears to be the case that there are oligomers larger than trimers present on the gel. The intensity plot for the wild-type Lpp appears easier to quantify (Figure S2a), and there is a clear gap in the integrated signal separating the monomer population from that of the oligomeric populations, but even here there are significant uncertainties in where to place baselines, how to assign certain regions on the gel to the dimer and trimer populations, and whether or not higher-order oligomers are also present. For these reasons we have considered a variety of ways of determining the relative populations of the various oligomeric forms (see Figure S3). The resulting relative populations vary, but a reasonable estimate is that Lpp chains that are in monomeric, dimeric, and trimeric forms are present in respective proportions of roughly 69:24:7.

With what we consider to be a semi-quantitative estimate of the relative populations, we wrote simulation code that explicitly models the cross-linking experiment under the same assumptions as those adopted by the Silhavy group (see Materials & Methods). The simulations use the stochastic simulation algorithm (SSA) developed by Gillespie and implemented using the so-called “direct method” (46,47); each step of the simulation involves the addition of a DSS-mediated cross-link between two Lpp chains within the same Lpp trimer. We performed simulations for a variety of scenarios ranging from one in which all Lpps are assumed to be resident in the periplasm to one in which two-thirds of the Lpps are assumed to be exposed on the cell surface (see below). A schematic illustrating how the cross-linking simulation proceeds is shown in Figure 2a. We start at the top of the Figure with an example scenario in which we assume that there are two Lpp trimers exposed on the cell surface for every ten Lpp trimers that are in the more conventional periplasmic location: covalent linkages between the periplasmic Lpps and the PG are indicated by short black lines connecting them. Importantly, it can be seen that the total number of Lpp chains in the schematic that are covalently connected to the PG is twelve, with eight of the periplasmic Lpps being connected once, and two of the periplasmic Lpps being connected twice (see Lpps at the right-hand side). Since there is a total of 36 Lpp chains in the simplified system shown, this means that one-third of the Lpp chains are covalently connected to the PG. This is a constraint that we enforce in all of the simulated scenarios in order to ensure consistency with the Inouye group’s early result that one-third of cellular Lpp chains are covalently connected to the PG (48). Throughout Figure 2a, Lpp chains are colored according to the way in which they would be observed in the experiment described by Cowles et al. (15): red, green, and blue cylinders represent respectively uncross-linked monomers, cross-linked dimers, and cross-linked trimers that should be observable in the experiment since they are not covalently connected to the PG. Yellow cylinders, on the other hand, indicate those Lpp chains that should not be observable in the experiment since they are covalently connected to the PG (directly or indirectly). In the example scenario shown in Figure 2a, therefore, 12 of the Lpp chains are already considered experimentally unobservable even as the simulation begins.

**Figure 2.**
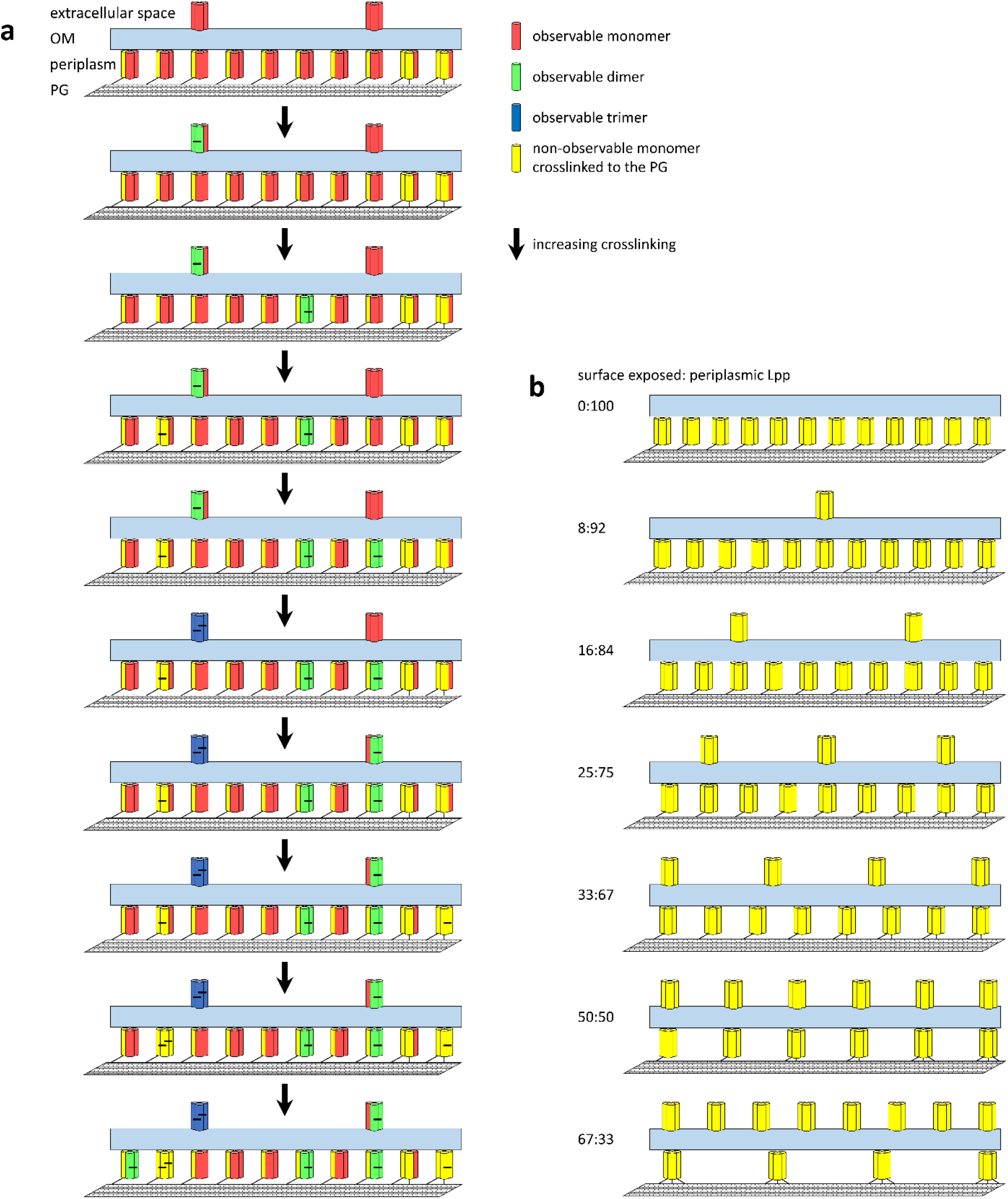
Scenarios considered in stochastic simulations of DSS-mediated cross-linking. (a) Illustration of one possible sequence of cross-linking events and their effects on the counts of experimentally observable and non-observable DSS cross-linked Lpp oligomers. (b). Illustration of different scenarios of partitioning of Lpp between periplasmic and surface-exposed subcellular locations; numbers at left indicate the percentages of Lpp trimers assigned to the cell surface and the periplasm, respectively. Note that the number of covalent connections between Lpp chains and the PG is identical in all scenarios.

The simulation proceeds stochastically in a series of timesteps, with each such step involving the addition of a single cross-link between pairs of lysine sidechains within a single Lpp trimer (see Materials & Methods for details). In the simplified example shown in Figure 2a, the first added cross-link is randomly selected to occur within one of the surface-exposed Lpps; addition of this cross-link (short black horizontal bar) has the effect of increasing by one the number of observable dimers and decreasing by two the number of observable monomers. The second added cross-link happens to be within a periplasmic Lpp, but the outcome is the same. The third added cross-link also involves a periplasmic Lpp, but in this case it cross-links a PG-connected Lpp chain to one that is unconnected; this has the effect of decreasing the number of observable monomers by one without increasing the numbers of observable dimers or trimers. The fifth added cross-link occurs to a surface-exposed Lpp, and since in this case it causes all three chains of the trimer to be covalently connected to each other, the system now contains an experimentally observable trimer. The simulation proceeds in this way, with the total numbers of observable monomers, dimers, and trimers being monitored continuously. Eventually, assuming that each periplasmic Lpp trimer contains at least one covalent connection to the PG (see Discussion), all of the periplasmic Lpp chains will be rendered unobservable as they become cross-linked to chains that are connected in some way to the PG. Importantly, throughout the simulation, care is taken to properly account for the ways in which cross-links are likely to be physically accommodated within the structure of the Lpp trimer solved by the Lu group (49): there are twelve lysine sidechains in total in an uncross-linked Lpp trimer, but only certain combinations of these sidechains provide plausible locations for DSS-mediated cross-links to occur (see Materials & Methods).

The simple scenario sketched in Figure 2a contains only 12 Lpp trimers. In actual simulations (which were performed in triplicate) we include 397,213 Lpp trimers, which is the copy number of Lpp estimated by the Weissman group (36), and we consider a wide variety of possible distributions of Lpps between the cell surface and the periplasm. Each of these situations is represented schematically in Figure 2b, again in a schematic form that shows only 12 Lpp trimers. All of the scenarios that we investigate are formulated in such a way that they remain consistent with the Inouye group’s finding that one-third of Lpp chains are covalently attached to the PG (48). At one extreme (top of Figure 2b), we consider a model in which 100% of the Lpps are periplasmic, i.e., the “textbook” view of Lpp distribution. In this case, the simplest way of ensuring that one-third of all Lpp chains is PG-connected is to assume that one of the three chains in every Lpp trimer is connected; it is also, however, to imagine a situation in which a major population of singly-connected Lpps is accompanied by equal minor populations of doubly-connected and unconnected Lpps (see Discussion). At the other extreme (bottom of Figure 2b), we consider a model in which two-thirds of the Lpp trimers are assumed to be surface-exposed: this is the view suggested by Cowles et al. in which the entire sub-population of “free” form Lpp is surface-exposed, and the entire sub-population of “bound” form Lpp is resident in the periplasm. We note that in this scenario, the only way to satisfy the Inouye group’s one-third PG-connected criterion is to assume that *every* chain of every periplasmic Lpp trimer is covalently connected to the PG. In between these two extremes lie a number of different intermediate models. The goal of the cross-linking simulations that we perform is to determine which, if any, of these models produces relative populations of monomers, dimers, and trimers that match the relative populations identified in the experiments of Cowles et al. that we quantified above.

An example time-course of one such simulation – that in which one-sixth (17%) of all Lpp trimers are considered to be surface-exposed – is shown in Figure 3a. At time zero, no cross-links are present, so the population consists solely of uncross-linked monomers. As time progresses, however, cross-links are added and the population of uncross-linked monomers decreases, either because they are converted into observable cross-linked dimers or because they become cross-linked to PG-connected chains and so drop out of the observable population (see above). Monomers encountering the first of these two fates lead to a rapid increase in the population of cross-linked dimers early in the simulations that is followed, after a noticeable lag, by a slower increase in the population of cross-linked trimers. Eventually – after a time much longer than the 20-minute time frame of the actual experiment (15) – trimers are predicted to dominate the observable population as lower-order oligomers are converted either into observable trimers or drop out of the observable population due to connection to the PG; by definition in the simulation, observable trimers can only originate from Lpps that are not covalently connected to the PG.

**Figure 3.**
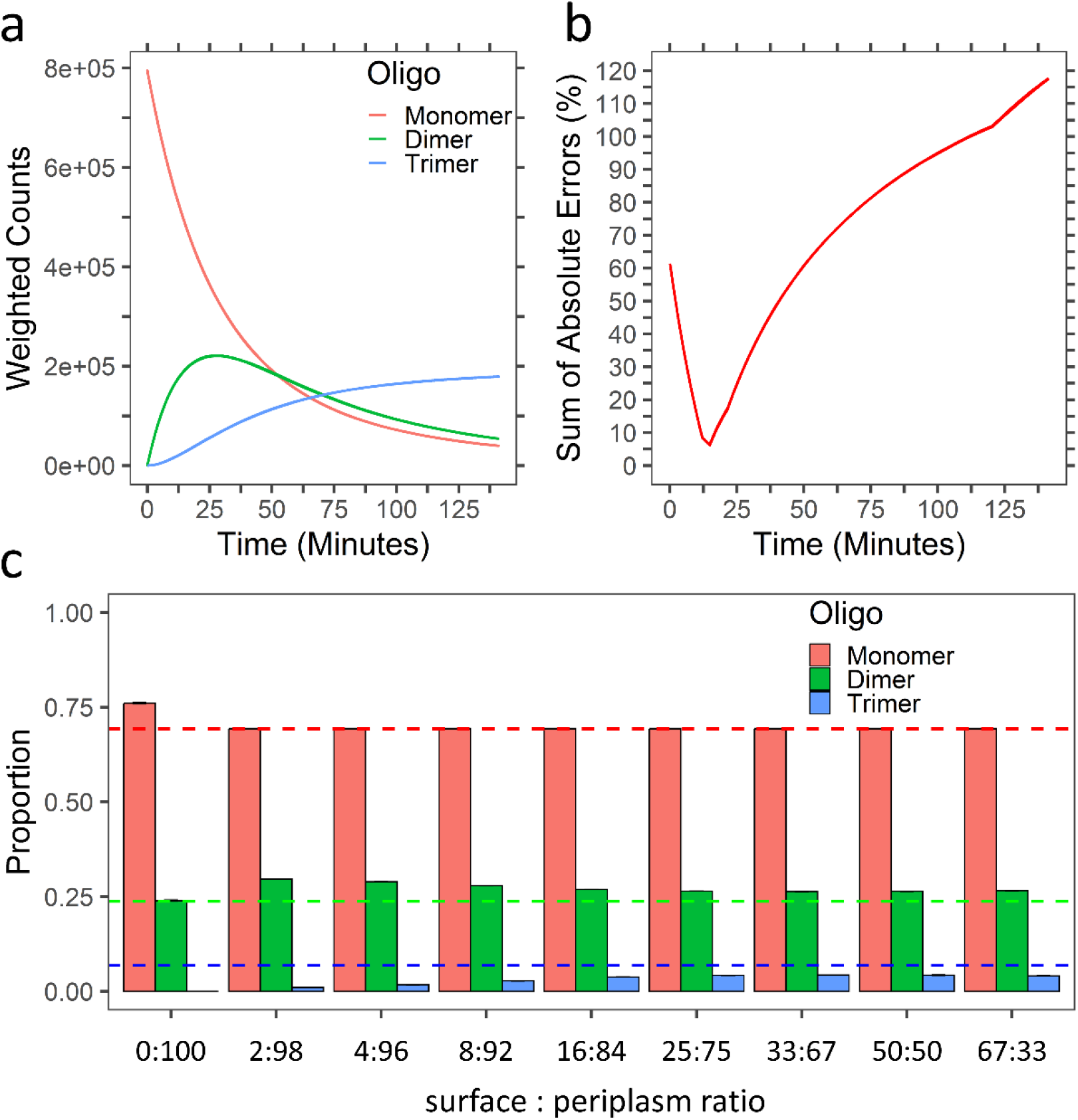
Results of stochastic simulations of DSS-mediated cross-linking. (a) The time course of a stochastic simulation performed in which 17% of the Lpps are considered to be surface-exposed. Lines show the weighted counts of DSS cross-linked Lpp monomers, dimers, and trimers colored in red, green, and blue, respectively. Counts are weighted according to the number of chains in each oligomer type. (b) The time course of the same reaction shown in (a) but plotting the sum of absolute errors between the simulated and experimental relative populations of all three types of DSS cross-linked oligomers. Note that the error is minimized at a time point roughly commensurate with the 20-minute timescale of the corresponding experiments (15). (c) Comparison of the optimal simulated relative populations of all three types of DSS cross-linked oligomers for all tested ratios of surface-to-periplasmic ratios. The dashed horizontal lines represent the experimentally determined relative populations with each colored according to the oligomer type that they represent. Note that the level of agreement is essentially constant for all surface-to-periplasm ratios greater than 17:83.

The absolute populations shown in Figure 3a can be converted into relative populations and compared with the corresponding experimental estimates that we noted earlier. A plot showing the combined absolute errors of these three relative populations versus simulation time is shown in Figure 3b, from which it can be seen that the error is minimized at a timepoint that roughly matches with the experimental timescale. An analysis identical to this was conducted for simulations of all tested Lpp distributions, and the relative populations obtained when each of these simulations produced their lowest errors are compared with the corresponding experimental estimates in Figure 3c. In all scenarios, the simulations somewhat overestimate the relative population of dimers while underestimating the relative populations of trimers, but the level of agreement with experiment is clearly worse for those distributions that place all or almost all of the Lpp within the periplasm. At a qualitative level, then, the simulations support the Silhavy group’s contention that at least some fraction of the Lpp must be exposed on the cell surface. However, once the fraction of Lpp that is surface-exposed rises to ∼17% (i.e., 1/6 of the population), all distributions achieve a similar level of agreement with the experimental estimates. Since the scenario in which only 1/6 of Lpp trimers is surface-exposed appears to be as good as any other scenario for matching the experimental estimates, the simulations also provide us with a view of the partitioning of Lpp that is much more in line with the conventional view that most Lpp resides in the periplasm. For the moment, then, it is this distribution with which we proceed here.

### A program for constructing whole-cell models of the peptidoglycan

The work presented up to this point serves principally to define critical input parameters required to guide the construction of whole-cell PG models. Coupling these data with: (a) experimental data on the populations of the various peptide sidechains and cross-link types, which here we get from Obermann and Höltje’s work (11), and with (b) an estimate of the likely distance between adjacent hoops of PG of 25 .5 Å, which is well within the range 27±5 Å indicated by AFM measurements (17) we are in possession of all the input parameters needed to build whole-cell PG models. Additional computer code was written, therefore, that uses these inputs to build structural models of the PG in which glycan chains are modeled at a resolution of 1 bead per sugar, and peptide sidechains and cross-links are modeled at a resolution of 1 bead per two amino acids. Full details of the model-building code are provided in Materials & Methods, but a central assumption of the code is to build glycan strands in an idealized arrangement in which all strands are oriented perpendicular to the long axis of the cell. In a further simplification we assume that these strands are laid down in a series of complete “hoops” arrayed from one end of the cell to the other, with all strands within a given hoop assumed to have the same clockwise or counterclockwise orientation relative to the long axis. The extent to which adjacent glycan strands are arranged parallel or antiparallel to each other within living *E. coli* cells does not yet appear to be established definitively, so we adopt a simple strategy of borrowing from sophisticated simulations reported by the Jensen group (29,30) which suggest that glycan strands may be added in pairs during both cell elongation and division. We therefore assume, for simplicity, that the orientations of strands within a series of successive hoops all follow the sequence …CCccCCcc… where C represents clockwise and c represents counterclockwise orientations. We justify this simplified approach on the grounds that the CG models that we build here are insufficiently detailed for the direction of glycan strand deposition to matter very much.

Having determined the relative strand orientations of each hoop of PG, the 3D model-building code, which we call *PG_maker*, randomly selects glycan strand lengths for placement within each hoop in such a way that the glycan length distributions derived earlier are automatically reproduced in both the polar and cylindrical regions of the cell (see Materials & Methods). Having placed all of the glycan strands, *PG_maker* then attempts to add each of the major chemical species that are found in the PG of *E. coli*. As such, the code allows the user to specify desired percentages of (a) seven different kinds of peptide sidechains, (b) three different kinds of dimeric peptide cross-links (tetra-DD-tetra, tetra-DD-tri, and tetra-LD-tri), (c) one type each of trimeric and tetrameric peptide cross-links, and (d) five different types of covalent connections between the PG and trimeric Lpp molecules. Importantly, different desired compositions can be defined for the polar and cylindrical regions of the cell. Draft whole-cell models can be generated with *PG_maker* in under an hour, and these generally match all desired populations of the various requested components to within 5%. The placements of certain components within the model, however, are subject to geometric constraints, e.g., MurNAc residues cannot be connected by peptide cross-links if they are more than 35 Å away from each other, or if their peptide sidechains are not oriented toward each other. It is not guaranteed, therefore, that the desired proportions of all components will be reproduced immediately. Since *PG_maker* automatically reports an inventory of all placed components, however, closer matches to the desired proportions of components of interest can be achieved by iteration, with the target values set in *PG_maker*’s input file being updated until the desired populations are achieved.

To illustrate *PG_maker*’s capabilities we focus here on building and analyzing a whole-cell PG model appropriate to an average sized *E. coli* cell in rich defined media (50), with Lpps assumed to be partitioned between the periplasm and the cell surface in the 5:1 ratio noted above. The final PG model produced in this way is shown in Figure 4a, which also displays a zoomed-in section of the cell; Figure 4b provides an illustration of some of the categories of components included. The final model contains 132,250 glycan strands with a total of 3,662,096 DS units, 1,478,055 free peptide sidechains, 907,633 million peptide cross-links of various types, and 307,543 thousand Lpp trimers. The effective surface area per DS unit that we measure from the model (240 Å^2^) is in good agreement with the experimental estimate of 250 Å^2^ (51). Perhaps more importantly, the final model’s inventory of components matches extremely well with the initial input specifications (Table 1): this indicates that the parameters and copy numbers of the various components derived from experiment and from the earlier simulations are all physically realizable in a single 3D structural model. Of the various components built into the model, the largest deviation between input and output populations (an error of 21%) is found with the tetrameric cross-links, which while rare, are difficult to place within the model since we require that they connect four separate glycan strands in only two adjacent PG hoops (see Materials & Methods).

**Figure 4.**
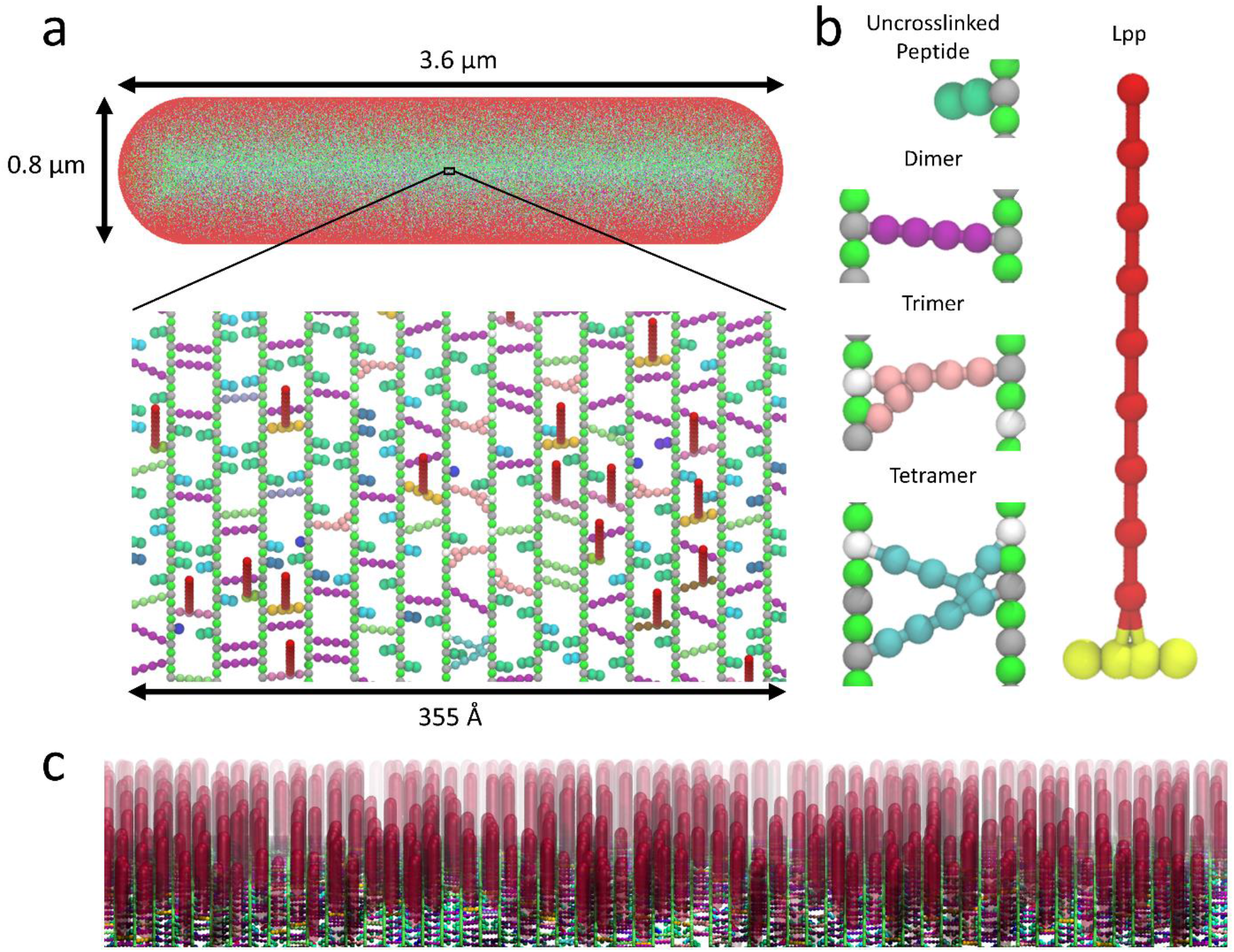
A whole-cell PG model built using *PG_maker*. (a) A visual representation of a PG model for an average sized *E. coli* cell growing in rich defined media. The zoomed-in image shows a small patch of the PG in CPK representation, with glycan strands illustrated such that green, grey, and white spheres represent GlcNAc, MurNAc, and 1,6-anhydro MurNAc residues, respectively. Different types of free (i.e., uncross-linked) peptide sidechains and cross-links are colored according to their identities; Lpp trimers are shown in red. (b) Illustration of some of the general component types built into the PG model shown in CPK representation. (c) A view showing the high density of Lpp trimers built into the model, protruding out of the PG layer and connected to the outer membrane (not shown). These images were generated with VMD (61).

**Table 1.** Summary of the counts of various PG components built into the PG model built by *PG_maker* and shown in Figure 4a. Also shown are the counts of the same components requested by *PG_maker’s* input file, together with the percent deviation between the two.

| Component Name | Actual | Expected | % Error |
| --- | --- | --- | --- |
| Dimer DD | 595759 | 595523 | 0.04 |
| Dimer DA | 92075 | 91556 | 0.57 |
| Dimer LD | 42261 | 41226 | 2.51 |
| Trimer | 86080 | 86122 | -0.05 |
| Tetramer | 1788 | 2277 | -21.49 |
| Lpp (Singly Linked) | 156627 | 156651 | -0.02 |
| Lpp (Doubly Linked) | 61246 | 61256 | -0.02 |
| Lpp Dimer DA | 76106 | 75426 | 0.90 |
| Lpp Dimer LD | 13564 | 12945 | 4.78 |
| Glycan Strands 1 – 5 DS units | 52298 | 51669 | 0.60 |
| Glycan Strands 6 – 10 DS units | 29327 | 29366 | -0.07 |
| Glycan Strands 11 – 20 DS units | 25406 | 25624 | -0.43 |
| Glycan Strands 21 – 30 DS units | 8432 | 8589 | -0.92 |
| Glycan Strands 31 – 50 DS units | 1153 | 1107 | 2.04 |
| Glycan Strands 51+ DS units | 15634 | 15720 | -0.27 |

One important independent check that we can place on the models concerns the predicted total cellular copy number of Lpp. *PG_maker* determines the number of Lpps to add by designating user-defined percentage of MurNAc residues as having attached Lpps which, in this case, is 12.5% (11). We obtain the predicted total cellular number of Lpps number by multiplying the number of Lpps covalently attached to the PG model by 1.2 to account for the 17% of Lpps that we assume are surface-exposed (see above). The resulting number – 369,051 – compares very well with the average Lpp copy number – 397,213 – estimated by the Weissman group on the basis of their ribosome-profiling experiments performed in the same media (36). Interestingly, the very high number of Lpps built into the model results in a highly crowded situation in the compartment of the periplasm that lies between the PG and the OM (Figure 4c). We explore the potential consequences of this crowding on the access of periplasmic proteins to this compartment in a subsequent section (see below).

### Preliminary estimates of pore size distribution in whole-cell models of the peptidoglycan

We anticipate that the 3D models produced by *PG_maker* will eventually serve as starting points for large-scale simulation studies (see Discussion), but even in their current, idealized form they allow us to conduct several preliminary analyses of their structural properties. The first property that we consider is the distribution of pore sizes within the PG since this has obvious implications for the ease with which periplasmic proteins might be able to travel back and forth between the inner- and outer-membranes. To quantify pore sizes we have used image processing software similar to that used by the Foster group (17) to analyze their AFM studies of PG sacculi (see Materials & Methods). Examples of the segmentations obtained from ImageJ (44) are shown in Figures 5a and 5b for the cylindrical and polar regions of the cell, respectively. From these images it can be seen that pores that result from the model-building process span a wide range of sizes: while there are many smaller pores, there are also some very large pores. A histogram of the pore sizes identified by ImageJ (quantified by their area) is shown in pink in Figure 5c; if the smallest histogram bin is omitted from consideration, the pore sizes in the PG model are exponentially distributed with a decay constant of −0.0027 Å^−2^. Figure 5c also shows (in green) corresponding data reproduced from the AFM imaging experiments of the Foster group (17); these data also fit to an exponential function, in this case with a decay constant of −0.0025 Å^−2^. Apart from the obvious difference between the computational and experimental distributions for the smallest pore sizes, therefore, the highly idealized PG model generated by *PG_maker* produces pore size distributions that are in surprisingly good agreement with experimental estimates.

**Figure 5.**
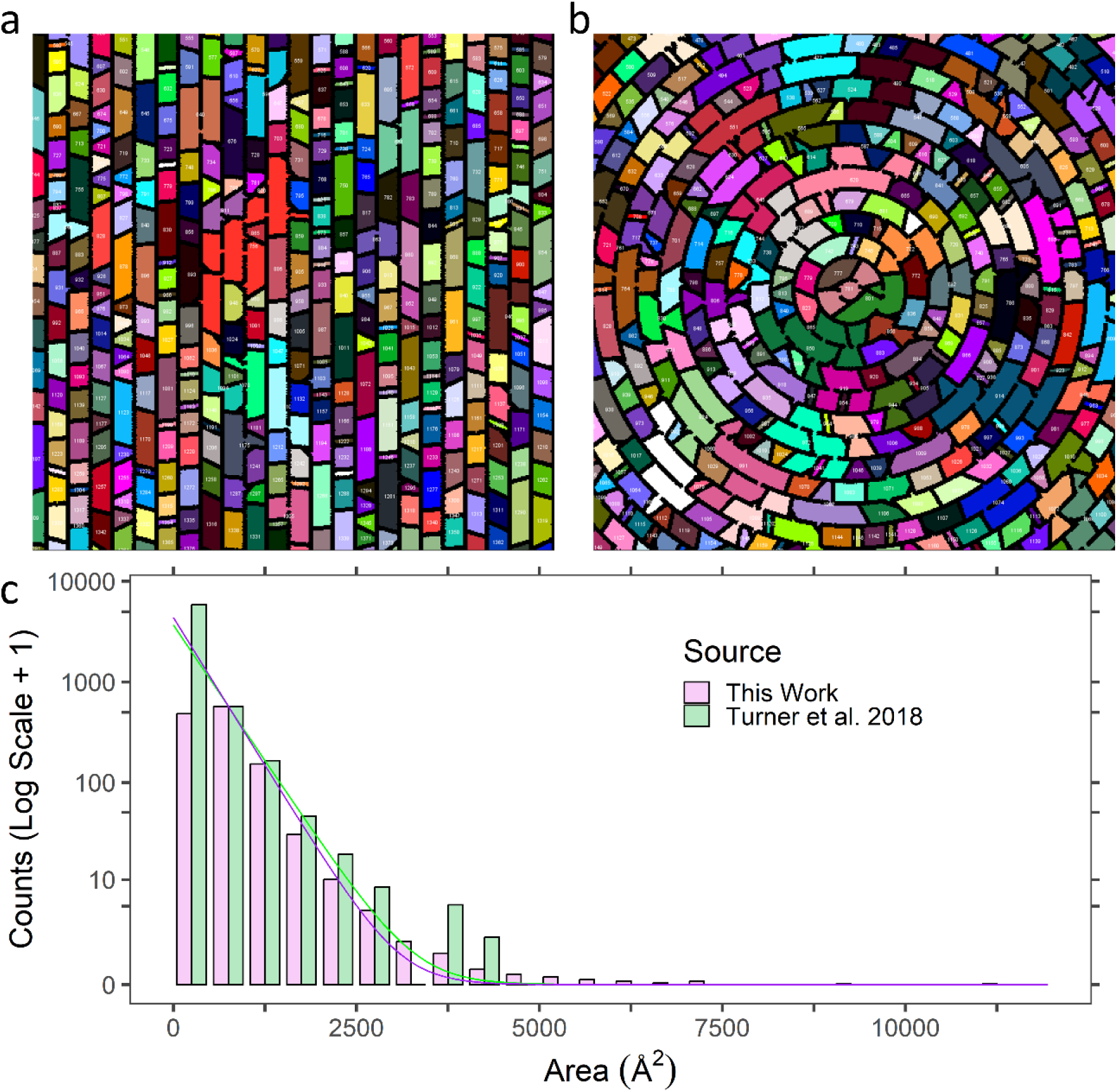
Pore size distributions in the 3D model of the PG produced by *PG_maker*. (a) and (b) show views of pores identified from images of the PG by ImageJ/Fiji (44,45) for the cylindrical and polar regions of the cell, respectively. (c) Distribution of the pore sizes identified from the PG model built in this work and from the data reported by the Foster group (17). The lines show the exponential decay functions fitted to each data set (see main text).

### The high periplasmic population of Lpp may limit access of periplasmic proteins to the OM

A second question that we have addressed with the structural model shown in Figure 4a is the extent to which the highly crowded “forest” of Lpps (Figure 4c) is likely to hinder access of proteins to the OM-proximal compartment of the periplasm. To explore this issue, we wrote additional computer code that computes a free energy profile characterizing how the thermodynamics of transferring a periplasmic protein from the IM to the OM is affected by steric clashes between the protein and the complete PG model. Full details of the calculations are provided in Materials & Methods, but in brief the calculations proceed by repeatedly placing each protein, in randomly selected orientations, at randomly selected positions around the cell and determining the extent to which their positions clash with either the IM or the OM or with the PG model. Calculations were performed on five proteins known to be resident in the periplasm of E. coli and exhibiting a range of sizes and shapes (Figure 5a). Figure 5b shows the free energy profiles calculated as a function of radial distance for the smallest of these five proteins (yahO; 70 residues once its signal peptide has been cleaved). Two lines are plotted. The first line (red) shows the free energy profile obtained when the PG is included in the calculations but when the Lpps are omitted. Reading from left to right, ΔG starts with a positive value owing to the fact that most of the randomly selected orientations of the protein are sufficiently close to the IM that they clash with it, ΔG then drops to zero as the protein reaches a radial distance at which none of the orientations clash with the IM, before it rises again as orientations begin to clash with the PG. As the protein attempts to pass through the PG layer ΔG reaches a maximum – in this case of ∼7 kcal/mol – reflecting the fact that there are very few orientations of the protein, and very few possible positions around the cell, where the protein can pass through entirely unscathed. Having passed through the PG, however, ΔG again drops to zero in the compartment proximal to the OM before rising again as orientations begin to be rejected due to clashes with the OM.

The second (blue) line shows the free energy profile obtained when both the PG and the LPPs are present. As expected, the profile for the first part of the periplasmic journey is unchanged since there are no Lpps present in the periplasmic compartment proximal to the IM. As the protein approaches the PG, however, the profiles begin to diverge: ΔG increases owing to the fact that some orientations that are non-clashing in the absence of the Lpp, are clashing in its presence. This has the effect of raising the free energy barrier associated with crossing through the PG slightly but has a much greater effect once the protein passes through the PG: in particular, in contrast to what was observed in the absence of Lpp, ΔG does not drop to zero. Instead, it plateaus at a value of ∼0.35 kcal/mol, which reflects the fact that a significant fraction of the randomly selected orientations of the protein clash with one or more Lpps. Overall, then, comparison of the two free energy profiles suggests that the inclusion of Lpp modestly increases the barrier to passing through the PG but has a more substantial effect on the free energy of the protein once it is there. Put simply, therefore, the crowded environment generated by the high density of Lpps in the compartment proximal to the OM makes it thermodynamically unfavorable (from a purely excluded-volume perspective) for the protein to enter this compartment relative to the IM-proximal compartment.

The resulting free energy profiles obtained for all five tested proteins in the presence of both PG and Lpp are shown in Figure 5c. At a qualitative level, all of the plots are identical: all of the proteins are excluded from the IM by steric interactions, all of them reach a ΔG of zero when they are sufficiently far from both the IM and the PG, all of them experience a large free energy barrier as they attempt to cross the PG, before the ΔG again drops as they reach the compartment that lies between the PG and the OM. At a quantitative level, however, the free energy profiles differ substantially, reflecting both the sizes and the shapes of the proteins. The three largest proteins in terms of residue counts (ptrA, livJ, and yncE) are sufficiently large that none of the 10 million paths successfully pass through the PG and we cap the ΔG at 9.4 kcal/mol as a lower-bound on the size of the free energy barrier. The smallest free energy barrier that we measure is for osmY, which at 173 residues is considerably larger in molecular weight terms than yahO. The lower computed free energy barrier for osmY reflects its elongated shape, which allows it to more easily negotiate its way through the PG’s pores. But this comes at the expense of its ability to fit into the OM-proximal compartment of the PG: in comparison with the larger proteins livJ (344 residues) and yncE (323 residues), it is more thermodynamically penalized for being in the OM-proximal compartment. This is because for radial distances at which livJ and yncE can comfortably find interstices between the PG, the OM, and the surrounding Lpps, osmY’s elongated shape hinders it from doing so. The largest protein investigated, ptrA, which at 939 residues is much larger than any of the others, experiences the highest thermodynamic penalty for accessing the OM-proximal compartment: even when it is positioned squarely in the middle of the OM-proximal compartment steric clashes with its surroundings are sufficiently prevalent for its ΔG to be elevated to 1.4 kcal/mol.

## Discussion

Here, we have attempted to show how a wide range of experimental data, coupled with a number of bespoke simulation codes developed during the course of this study, can be combined to produce self-consistent 3D structural models of *E. coli*’s PG on a whole-cell scale. Along the way, we have proposed what we think is a plausible distribution of glycan strand lengths that appears to be consistent with all the available experimental data, and we have developed a stochastic simulation code that explicitly models a critical earlier Lpp cross-linking experiment. The results obtained with the latter code may be particularly important as they suggest a potential “middle ground” resolution of a long-standing question concerning the spatial distribution of Lpps within the cell (see below). Having developed a second code, *PG_maker*, to construct 3D models of the PG, we have provided two examples of realistic emergent properties that arise naturally from their construction. First, we have shown that the copy number for Lpp predicted for the PG model of *E. coli* cells grown in rich defined media is in excellent agreement with the copy number estimated experimentally for the same growth conditions by the Weissman group (36). Second, we have shown that the distribution of pore sizes within the same model matches surprisingly well with corresponding AFM estimates reported by the Foster group (17). Finally, we have provided an example of the potential utility of whole-cell models for hypothesis generation: we have shown how the free energy of passage of proteins across the periplasm can be strongly influenced by the protein’s size and shape and by the presence of large numbers of Lpps in the periplasmic compartment that lies between the PG and the OM. In what follows, we attempt to place some of these results in context, discuss some of the limitations of the approach, and identify some possible future directions.

In Results, we began by addressing two quite different problems that have a potentially crucial bearing on the construction of whole-cell PG models. For the glycan strand length distributions, we showed using simple algebra that the available experimental data demand the existence of a substantial population of very long strands; the existence of such a population is entirely consistent with the Foster group’s AFM studies identifying glycan strands with lengths approaching 200 DS units (16). To develop a model capable of simultaneously describing all the known data regarding glycan lengths we assumed here that the length distribution is bimodal, with the previously uncharacterized distribution of the long-length component having a simple Gaussian distribution with an arbitrarily defined spread, and a mean value optimized to match experiment. We have shown that altering the width of the Gaussian has little effect on the derived center of the distribution (Figure S1), but beyond this all that we can really say is that our assumed model is not unreasonable since it is at least qualitatively consistent with the processive action of elongasomes on mreB filaments that are responsible for adding most of the PG in the cylindrical regions of the cell (40,52). There are, of course, likely to be other ways to fit the experimental data, but we think that all successful distributions are likely to be at least bimodal in form. Obviously, it would be most helpful if the glycan length distribution could be resolved experimentally over a wider range of lengths than has been reported previously. If available, experimentally determined length distributions could be passed immediately as input to *PG_maker* to produce structural models (see below).

The second major problem that we attempted to address prior to undertaking 3D model building efforts concerned the vexing question of the subcellular distribution of Lpp. At the outset, we hoped that by explicitly simulating the interesting cross-linking experiment conducted by Cowles et al. we might be able to determine, more or less exactly, the relative populations of periplasmic and surface-exposed Lpps. In practice, however, our ability to extract solid quantitative estimates of the relative populations of uncross-linked monomers, DSS cross-linked dimers, and DSS cross-linked trimers in the experimental data is sufficiently limited that we should probably consider the derived numbers to be only semi-quantitative estimates. But even putting that issue to one side, it appears that there is a wide range of possible scenarios for partitioning Lpp between the periplasm and the cellular surface that are likely to be equally capable of matching the experimental data. Our preference here has been to select a model in which ∼17% of the Lpps are surface-exposed, since this appears to provide a reasonable compromise in terms of building PG models that are generally consistent with the more “conventional” view of Lpp’s subcellular location, while simultaneously acknowledging the experimental data indicating that a significant fraction of cellular Lpp is exposed on the cell surface (15). It should be noted that the Silhavy group’s evidence in favor of a surface-exposed population is not limited to the single cross-linking experiment that has been our focus here, and work reported by the Brissette group has provided additional support (53).

We should also acknowledge that despite our efforts to build a realistic simulation model of the pivotal cross-linking experiment described in Cowles et al. (15), none of the tested distributions produces simulation results that match the experimental data especially well (see Figure 3c). It is important to note that this could be due to a wide variety of reasons. First, as noted above, our ability to accurately quantify the experimental data is limited, and this is especially problematic given that there appear to be subpopulations of species within the gel that defy easy categorization; this appears to be true especially for the ΔK_58_ Lpp mutant data (Figure S2b). It seems likely that some of the density on the gel represents alternative cross-linked dimers and trimers whose mobility differs due to the positions of their cross-links along the Lpp chain (see Materials & Methods). But there are also hints of the presence of one or more higher-order oligomers on the gel, with perhaps the most likely culprit being a DSS cross-linked tetrameric form of Lpp. One possible source for such a cross-linked tetramer would be an intermolecular cross-linking event that is not accounted for in the simulations, i.e., one occurring between two native Lpp trimers, each singly-connected to the PG, that are close to each other in space. The high density of PG-connected Lpps in the 3D models generated by *PG_maker* suggests that intermolecular cross-linking events are a distinct possibility: visual inspection of the small patch of PG highlighted in Figure 4a, for example, identifies several pairs of Lpp trimers that could easily be bridged by DSS molecules. A second source of uncertainty concerns the relative reactivities that we assign to the lysine sidechains in the simulations. Here we have assumed that all such sidechains have equal reactivities, but one could easily imagine that they might differ in reality. For example, the K58 sidechains at the C-terminus could well be more reactive given the apparent flexibility of this residue: it is unresolved in any of the available crystal structures (see, for example, RCSB ID: 1EQ7). On the other hand, it is possible that the sidechains of surface-exposed Lpps might be relatively less reactive, especially those nearer the N-terminus, since they are likely to be at least partly masked by the high density of surrounding lipopolysaccharide molecules.

Finally, it is worth noting that while our currently preferred model is one in which ∼17% of the Lpps are surface-exposed, it is technically possible for a model that has no Lpps exposed on the surface to match the experimental cross-linking data with the same level of success as other models. All that is required for this to be true is that a fraction of the periplasmic Lpps be covalently unconnected to the PG; scenarios that achieve this while simultaneously reproducing the Inouye group’s one-third PG-connected criterion (48) can be constructed by requiring only that the major population of singly-connected Lpps is accompanied by minor but equal populations of unconnected and doubly-connected Lpps. In Figure S4 we show that assuming that as little as 17 % of the periplasmic Lpps are unconnected to the PG produces a match with the experimental cross-linking data that approaches that achieved with substantial populations of surface-exposed Lpps. The existence of a small population of unconnected Lpps within the periplasm is not unreasonable given the recent demonstrations that the periplasmic enzyme LdtF (previously named YafK and now renamed DpaA) catalyzes the hydrolysis of the covalent connection between the Lpp and the PG (54,55).

The best way to settle the subcellular distribution of Lpp in truly quantitative terms is obviously, in principle, via experimental techniques capable of probing living cells. If this can be done, then as was the case with the glycan strand length distributions, the updated data can again be input directly to *PG_maker* to produce corresponding 3D models. The *PG_maker* code is agnostic regarding the level of realism in the populations of the various chemical components requested by the user: put simply, its job is to build what is asked of it, and so long as the requested components are physically realizable in a 3D model, *PG_maker* will rapidly build it. The last point is important: the placements of many components are subject to geometric constraints that are intended to ensure the final models make physical sense. We can already identify two components for which these constraints are sufficiently restrictive that it can be challenging to find suitable positions to place them. The first is the tetrameric peptide cross-link (generally denoted tetra-DD-tetra-DD-tetra-DD-tetra) for which the code currently assumes that each of the cross-linked peptide sidechains must originate from a different glycan strand. This constraint allows us to build complex situations in which four different glycan strands are covalently inter-connected by a single nexus of cross-linked peptide sidechains, but it is also likely to be the reason why our whole-cell model slightly undercounts the requested numbers of tetrameric peptide cross-links in the whole-cell model (see Table 1). The second is the Lpp trimer that is triply connected to the PG, i.e., in which all three of the chains in the Lpp trimer are covalently connected to the PG: again, the code currently assumes that each of these chains should be connected to a different glycan strand. Attempts to build a whole-cell PG model along the lines suggested by the Silhavy group, in which every chain of every periplasmic Lpp trimer must be covalently connected to the PG, will therefore likely require relaxing some of the geometric assumptions on which the *PG_maker* code currently relies.

The coarse-grained models that *PG_maker* constructs fall some way short of providing an atomic representation of the PG, but they are significantly more detailed than has been used in large-scale CG simulations in the past (26–30). Importantly, we think that the current resolution of the models provides a good compromise between the requirement for as few beads as possible, in order to eventually make it feasible to perform future dynamic simulations of the PG on a whole-cell scale, while retaining a sufficient number to realistically describe the PG’s conformational properties. In particular, the approach we have adopted affords glycan strands a considerable degree of flexibility, and the use of explicit CG beads to model the peptide sidechains and cross-links allows us to make estimates of pore sizes that appear to be in quite reasonable agreement with experiment (Figure 5c). Given the idealized nature of the structures built by *PG_maker* these comparisons should be considered somewhat preliminary, but it is striking that the only glaring anomaly that we obtain currently is that the frequency of occurrence of the very smallest pores – which are too small for any globular protein to pass through them – appears to be underestimated relative to experiment. It will be interesting to determine the extent to which these pore size distributions change as and when the PG model is subjected to dynamics simulations that allow it the freedom to sample alternative conformations. Similar considerations apply to our calculations of the thermodynamic profile for passage of proteins across the periplasm (Figure 6c). While the computed free energy profiles will certainly change at a quantitative level when less idealized PG structures are used in the calculations, we expect that the key results reported here will remain qualitatively unchanged: the ability of proteins to pass through the PG’s pores will remain a function of their shape, and the thermodynamic penalty that they pay for access to the OM-proximal compartment of the periplasm will continue to be dependent on the local density of volume-excluding Lpps. It should also be remembered, however, that the present calculations consider only steric interactions between proteins and the PG and Lpp. We know from our previous work, however, that the thermodynamics of proteins in intracellular environments are also subject to modulation by electrostatic and hydrophobic interactions (56), and the atomistic MD simulations reported by the Gumbart (20) and Khalid groups (24,25) have already demonstrated the potential for favorable interactions to occur between proteins and elements of the PG.

**Figure 6.**
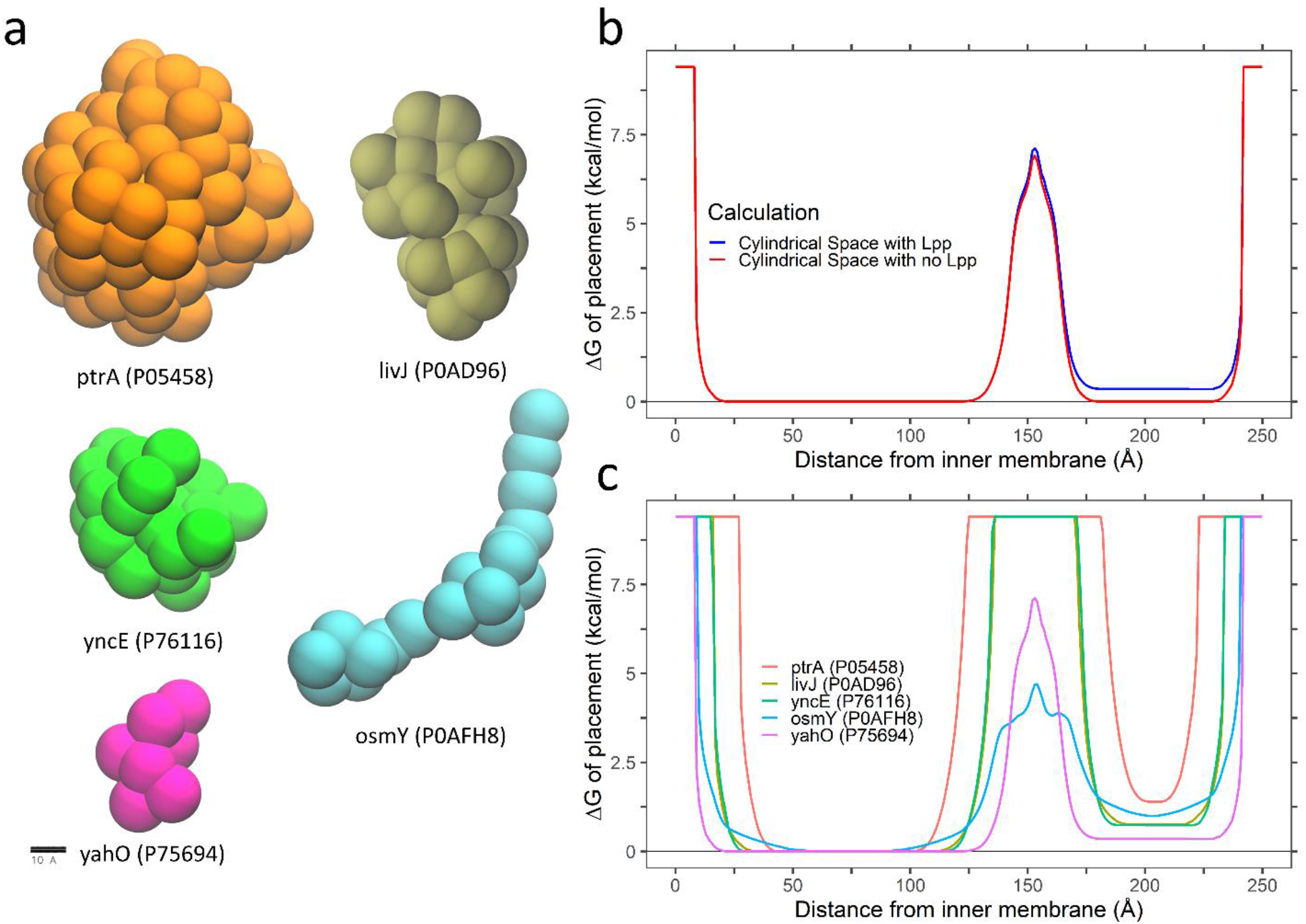
Excluded-volume dependent free energy profiles for proteins passing through the PG. (a) The five proteins for which free energy calculations were performed are shown in their CG representations. (b) Plot of free energy versus radial distance from the inner membrane for the small protein YahO in the cylindrical regions of the cell; separate lines are shown for calculations that excluded (red) or included (blue) Lpp in the calculations. (c) Plot of free energy versus radial distance for all five proteins with Lpp included in the calculations.

Finally, we point out some possible future directions for further development. The *PG_maker* code already allows a high degree of chemical and compositional heterogeneity to be incorporated into the 3D models. In particular, by allowing very different compositions to be assigned to the polar and cylindrical regions of the cell, it enables the models to reflect the fact that entirely different cellular machineries are used to build the PG in these two regions of the cell (57). In some respects, then, the code simply awaits the arrival of additional experimental data that can be used to further constrain the target compositions of the models. But other developments can certainly be imagined. One obvious direction would be to increase the resolution of the models, perhaps even to the atomic level; at such a resolution, the question of how likely adjacent glycan strands are to be oriented in the same direction will become more important to address (see above). A second obvious direction is to attempt to move beyond the highly idealized, single-layer structures that are currently output by the code. The most straightforward way to do this is to subject the *PG_maker*’s models to dynamic simulations, possibly performed on the whole-cell scale: such simulations would soon rid the models of their current highly regular, lattice-like appearances. This is a direction that we will be actively pursuing in the future. It may also, however, be possible to devise other ways of laying down glycan strands that do not depend on the uniformly spaced “hoops” that the code currently draws from. In particular, more irregularly added glycan strands would allow the models to reflect that fact that cellular PG is continually remodeled and repaired by machinery such as PBP1B and LpoB (57,58) and would likely produce localized regions where the PG may take on a double-layer appearance (19).

## Materials and Methods

### Deriving glycan strand length distributions for polar and cylindrical regions of the cell

The primary source of data that we use here to determine glycan strand length distributions is the work of Obermann and Höltje (11). Obermann and Höltje report the percentage abundances of glycan strands ranging in length from 1 to 30 disaccharide (DS) measured using high performance liquid chromatography (HPLC) of glycans containing radiolabeled N-acetylglucosamines; these same data have previously been used by the Wingreen group to construct large-scale PG models (26). Separate results were provided by Obermann and Höltje for: (a) so-called “minicells”, which are assumed by those authors to be completely composed of polar PG, and (b) whole cells, which they assumed in their subsequent analysis to be composed of a 2:7 ratio of polar and cylindrical PG. Since the detailed glycan strand length distributions are not provided in tabular form by Obermann and Höltje (11), we digitized Figure 3 from their paper and derived the values numerically using WebPlotDigitizer (https://automeris.io/WebPlotDigitizer/). In addition to providing detailed distributions for glycans of lengths 30 DS units and shorter, Obermann and Höltje reported the combined percentage of all glycans of longer lengths: their reported values for these combined populations are 8 and 13 % for minicells and whole cells, respectively. Finally, Obermann and Höltje also report the mean lengths of the glycan strands as 23.5 and 27.8 for minicells and whole cells, respectively. Importantly, the latter estimates were determined independently of the glycan length distributions by radiolabeling and quantifying the terminal anyhdro residues rather than the N-acetylglucosamine residues. To our knowledge, the Obermann and Höltje study is currently the sole available source of data that attempts to separately quantify glycan length distributions in the polar and cylindrical regions of the cell.

In order to develop models for the glycan length distribution over the entire range of possible lengths – which, to our knowledge has not been included in large-scale PG models previously – we proceeded as follows. The Obermann and Höltje length distributions for minicells and wild-type cells are very similar to each other from 5 to 30 DS units, and they both have what appears to be a simple power law decay. To provide a simple way of extrapolating these distributions to longer lengths, therefore, we decided to average them and fit the resulting data to a simple function of the form y = a x^−b^ + c. The resulting fit has an r^2^ value of 0.994 and produces the following parameter values: a = 0.2704, b = 0.6959, c = −0.0226. Since the fitted function passes through zero at 37 DS units it does not provide a meaningful description of the distribution for longer length strands. Following the reasoning outlined in Results, however, we know from simple algebra that there must be a population of glycan strands that are much longer than the 30 DS limit directly probed by Obermann and Höltje. To model the length distribution of these much longer strands we postulate that they follow a Gaussian distribution, and we further assume that the Gaussian has a standard deviation of 30 DS units. We also know from Obermann and Höltje that the combined contribution of all strands longer than 30 DS units must amount to 8 % of the total population for the minicells that we assume represent the polar PG. All that remains to be determined for the polar PG, therefore, is to find the optimal mean position of the Gaussian that describes the long strand population. This we do by trying all possible values from 38 to 400 DS units in intervals of 0.1 DS units and seeking the value that, when combined with: (a) Obermann and Höltje’s original data for 1 to 30 DS units, and (b) our power law fit for 31 to 37 DS units, results in the best match to the known mean strand length of 23.5 DS units (see above).

We follow a similar approach to deriving a glycan length distribution for the cylindrical region of the cell. But first, we must obtain an estimate of underlying length distribution for the glycan strands in the cylindrical region of the cell by combining Obermann and Höltje’s data for minicells with their data for wild-type cells. To do this, we follow Obermann and Höltje in estimating that 2/9 of the PG in whole cells is polar PG while the remaining 7/9 of the PG is cylindrical PG; this estimated partitioning was based on surface measurements of isolated murein sacculi (59). Using these estimates, we subtracted a suitably weighted version of the minicell (polar) length distribution from the wild-type (whole cell) length distribution to yield the estimated length distribution for the cylindrical PG from 1 to 30 DS units. We then obtained an optimal mean value for a long-strand Gaussian distribution using the same procedure described above for the polar PG (see above).

### Determining the abundances of peptide components required as input to *PG_maker*

The percentages of MurNAc residues connected to a wide variety of types of peptide sidechains and cross-links are reported directly in Table 1 of Obermann and Höltje (11); their Table 2 sums a number of these components into groups, apparently also adding on contributions from less abundant components that are not explicitly listed in their Table 1. Abundances are explicitly listed for six different types of free peptide sidechain, and all of these are included as separate components in the 3D PG models that we ultimately construct with *PG_maker* (see below). Abundances are also provided for 18 different types of dimeric peptide cross-link, 12 different types of trimeric peptide cross-link, and 2 different types of tetrameric peptide cross-link. Each of the dimeric cross-links listed can be considered a variant of one of the following three more abundant components: tetra-DD-tetra, tetra-DD-tri, and tetra-LD-tri. To keep the number of component types that *PG_maker* must be able to model manageable, the code is currently restricted to constructing only these three types of dimeric cross-links. For similar reasons, we assume that each of the trimeric and tetrameric cross-links can be considered variants of the more abundant tetra-DD-tetra-DD-tetra and tetra-DD-tetra-DD-tetra-DD-tetra components, respectively. Prior to constructing whole-cell PG models with *PG_maker*, the abundances of all of these components are scaled up by an average of ∼8% to reflect the contributions of minor components that were insufficiently abundant to be listed separately in Obermann and Höltje’s Table 1 but that were apparently included when the total amounts of component groups were listed in their Table 2. For the monomeric peptide sidechains in whole cells, for example, the sum listed in Table 2 is 7.2% larger than the sum of the individual components listed in Table 1; accordingly, we increase the required amounts of each free peptide sidechain type by 7.2%.

Obermann and Höltje’s Table 1 also lists the abundances of several components that contain a triLysArg species that is indicative of a covalent connection to an Lpp chain. To again keep the number of component types that need to be modeled by *PG_maker* manageable, the code currently allows Lpp trimers to be covalently connected to the PG via the following three mechanisms: (1) to one or more free tetrapeptide sidechains (see below), (2) to a dimeric peptide cross-link of the form tetra-DD-triLysArg, and (3) to a dimeric peptide cross-link of the form tri-LD-triLysArg. Each of the triLysArg-containing components listed in Obermann and Höltje’s Table 1 is approximated by one of these three cases, with their abundances again all scaled up by ∼8% to account for very minor abundance components that are not explicitly listed. Since *PG_maker* has the ability to place Lpp trimers so that they possess one, two or even three covalent connections to free peptide sidechains (each of which we assume must be part of a different glycan chain) we also need estimates of the ways in which free peptide sidechains containing a triLysArg component are partitioned between Lpp trimers containing one, two or three covalent connections to the PG. We obtain the latter from the results of our stochastic simulations of the DSS cross-linking experiment described by Cowles et al. (15) (see below).

*PG_maker* allows all components to be assigned different abundances in the polar and cylindrical regions of the cell. As noted above, we assume that the abundances of components in minicells reported by Obermann and Höltje provide direct estimates of their abundances in the polar regions (P_pol_) of wild-type cells. To derive the abundance of each component expected in the cylindrical regions of the cell (P_cyl_) we followed the assumption made by Obermann and Höltje that 2/9 of the PG is contained within the polar regions with the remaining 7/9 contained within the cylindrical region. Given the listed abundance of a particular component in whole cells (P_wc_) and in minicells (P_pol_), therefore, we can obtain an estimate of P_cyl_ using: P_cyl_ = 9/7 × (P_wc_ – (2/9) × P_pol_).

### Computer modeling of the chemical cross-linking experiment of Cowles et al

#### a. Quantifying relative proportions of chemically cross-linked Lpp

To determine the relative proportions of uncross-linked monomers, DSS cross-linked dimers and trimers in the immunoblots shown in Figure 6 of Cowles et al. (15), we used ImageJ/Fiji (44,45). Following a recently described protocol (60), a rectangle was drawn tightly around each lane using the “rectangle tool”, and the integrated intensity as a function of position along the gel was plotted using “plots lanes”. The resulting intensities are plotted in Figures S2a and S2b for experiments using wild-type Lpp and a ΔK_58_ mutant, respectively. Base lines and positional cutoffs were applied using the “straight line tool”, and the area under each peak was measured using the “wand tool”. The relative populations of the Lpp chains in monomeric, dimeric and trimeric forms were then obtained from direct comparisons of the areas under their corresponding peaks. As shown in Figure S3, a variety of extreme ways of analyzing the somewhat noisy signal were investigated in order to estimate potential uncertainties in the relative populations.

#### b. Stochastic simulation of the DSS cross-linking experiment

To mimic the DSS cross-linking experiment described by the Silhavy group *in silico*, we developed stochastic simulation code implementing the “direct method” algorithm formulated by Gillespie (46,47). Briefly, the code works as follows. The simulation proceeds in a series of steps, each of which involves the placement of a DSS-mediated crosslink between a pair of lysine sidechains contained within a single, randomly selected Lpp trimer. In reality, the cross-linking reaction almost certainly proceeds in at least two stages, with a DSS molecule first attaching to one lysine sidechain before subsequently reacting again with a second lysine sidechain nearby. Here, however, we model the process as a concerted and effectively bimolecular reaction dependent on the current populations of DSS molecules and reactive pairs of lysine sidechains (see below). The simulation contains 397,213 Lpp trimers (this being the total Lpp trimer copy number estimated by the Weissman group (36)), and an arbitrarily high number of DSS molecules (set here at 100 million) that ensures the latter remain in excess. Following exactly the approach outlined by Gillespie, at each step of the simulation the total reaction propensity, p_rxn_, is calculated as p_rxn_ = k_rxn_ × n_DSS_ × n_KK_, where n_DSS_ is the number of unreacted DSS molecules remaining, n_KK_ is the total number of lysine sidechain pairs that are physically capable of being bridged by a DSS molecule, and k_rxn_ is an effective bimolecular reaction rate constant that is scaled in such a way that the overall time course of the simulated reaction matches roughly with the experiment. In all the simulations described here k_rxn_ is set to 1 × 10^−13^. With the combined reaction propensity calculated, a uniformly distributed random number, u, in the range 0 to 1 is selected, and the time to the next reaction event, t_rxn_, is computed as t_rxn_ = ln (1/u) / p_rxn_. One of the cross-linkable lysine sidechain pairs is then randomly selected as the site of this next reaction, and the effects of this cross-linking reaction on the numbers of monomers, dimers, and trimers that would be observable in the experiment performed by Cowles et al. (15) are calculated as described in the main text.

An important point to note is that while the stochastic simulation framework formally considers only the numbers of the available reactive species, we can use knowledge of Lpp’s atomic structure to ensure that these numbers reflect the realities of the molecules involved. DSS is understood to be a cross-linker of primary amines with an ability to span a distance of ∼11 Å. In native Lpp trimers, there are five candidate sidechains that are likely to be capable of engaging in cross-linking by DSS; these are K6, K20, K39, K55, and K58 (residues numbered here assuming that the tripalmitoylated cysteine is number 1). We have built a model of the complete Lpp trimer using in-house homology code that adds in the C-terminal lysine that is missing from the Lpp crystal structure (49) (RCSB ID: 1EQ7; Figure S5). In it, we find that the lysine sidechains appear to form four separate “rings” along the long axis of the molecule that are all well separated from each other. The first three rings contain three lysine sidechains, while the fourth contains six lysine sidechains (those of K55 and K58 from each of the three chains). Based on our model of the full atomic structure of the Lpp trimer, therefore, we assume in the stochastic simulations that cross-linking reactions can occur only between lysine sidechains that are part of the same ring. An Lpp trimer that has no covalent connections to the PG will therefore contain a total of 24 cross-linkable lysine pairs: there will be 3 cross-linkable pairs in each of the first three rings (e.g. the first ring contains K6(chain A):K6(chain B), K6(A):K6(C), K6(B):K6(C) as cross-linkable pairs), and there will be an additional 15 cross-linkable pairs in the fourth ring (since there are 15 ways to connect the combined 6 copies of K55 and K58 with a single cross-link). In contrast, Lpp trimers that contain one or two covalent connections to the PG will contain only 19 and 15 cross-linkable lysine pairs, respectively. This is because each covalent connection to the PG eliminates a K58 sidechain from the pool of cross-linkable sites, so if, for example, K58(A) is the site of attachment to the PG, then there is no longer the possibility of forming DSS-mediated cross-links at K58(A):K55(A), K58(A):K55(B), K58(A):K55(C), K58(A):K58(B) or K58(A):K58(C). Simulations of the ΔK58 Lpp mutant are performed in the same way as described above, with the exception that the total number of cross-linkable lysine pairs is now 12 since the fourth ring of the Lpp contains only K55 sidechains.

### Constructing 3D whole-cell PG models with *PG_maker*

The ultimate objective of the preceding sections is to establish key input parameters for the *PG_maker* code that we have developed to build 3D models of the PG on the whole-cell scale. *PG_maker*’s primary purpose is to assemble all of the known components of the PG, including the many thousands of copies of covalently connected Lpps, in ways that make physical sense. A secondary purpose is to do so without incurring egregious steric clashes that would likely prevent the resulting models from being used as starting points for future large-scale dynamics simulations. *PG_maker* is a stand-alone Fortran program that has no external dependencies and so can be compiled very easily (see Code Availability).

*PG_maker* begins by reading an input file provided by the user. This file specifies the user’s desired percentage abundances of a variety of PG components, with different target values being allowed for the polar and cylindrical regions of the cell, together with desired geometric parameters such as the length and radius of the desired model, and the desired spacing between adjacent glycan strands along the long-axis of the cell. All PG modes constructed here were 3.6 µm in length, with the central 2.8 µm of this being the cylindrical region of the cell and capped at both ends by hemispheres, each with a radius of 0.4 µm. These dimensions are appropriate to average sized *E. coli* cells growing in rich defined media (50).

Once all input parameters have been read, the code starts by placing a series of “hoops” of glycan strands around the long axis of the cell. The separation distance between hoops (measured along the edge of the model) used in all models built here was set at 25.5 Å and the distance between adjacent sugars within a glycan strand was set at 4.70 Å; the latter number was obtained from exploratory molecular dynamics simulations of small peptidoglycan fragments. Treating each hoop independently, a randomly selected position around the hoop is chosen as a starting point, and a glycan chain length is sampled randomly according to separate probability distributions that are read in at run-time for the polar and cylindrical regions of the cell; these probability distributions reflect the glycan length distributions derived in the main text. If the selected glycan chain can be added without going past the hoop’s starting point, then it is added; otherwise, it is truncated at the number of glycans required to close the hoop. Truncating the chains in this way means that each hoop contains the exact number of glycans required to close it without leaving gaps or having glycans lie on top of each other, but it also has the undesirable effect of changing the final glycan length distribution in the 3D model so that it differs from the input probability distribution.

To correct this effect, we make the entire process of adding the glycan strands an iterative one, adjusting the input probability distributions until they produce the desired output. For a given type of PG model this iterative procedure only needs to be carried out once; additional replicate PG models can be made using the same adjusted probability distributions and they will automatically differ from each other owing to the inherently stochastic nature with which glycan strands are initially placed (see above). Figure S6 shows how the total error in the desired and output glycan strand length distributions becomes minimized during the iterative optimization process. Figure S7 shows how the final refined input strand length distribution differs from the actual desired distribution: note that the optimization process suppresses the probability with which very short glycan strands are selected and amplifies the probability with which very long strands are selected in order to ensure that the desired distributions are achieved.

Once glycan strands have been successfully placed around all hoops, the next stage of the construction process is to determine where to place peptide cross-links between glycan strands in adjacent hoops. An input parameter that becomes important at this stage is the periodicity with which peptide sidechains are assumed to point outwards from the glycan strands. In a model with a periodicity of 2 the directions of the peptide sidechains simply alternate: if the sidechain of one MurNAc points to the left, for example, then the sidechain of the next MurNAc in the strand will point to the right, and so on. In all of the models presented here, however, we have assumed a periodicity of 4: in a sequence of 4 DS units, two consecutive sidechains will point to the left (and therefore be available for cross-linking to glycan strands in the adjacent left-hand hoop), then the next two sidechains will point to the right (and be available for cross-linking to glycan strands in the adjacent right-hand hoop), etc. The starting point for the sequence of sidechain orientations in each glycan strand is left deliberately undefined until their first peptide cross-link is formed: once this happens, the directionality of all other sidechains is automatically determined given the input periodicity.

The first peptide cross-links whose placements are determined by *PG_maker* are the trimeric and tetrameric cross-links since these are the most difficult for which to find suitable placements. This is largely because we assume that trimeric cross-links must connect three different glycan strands (two of which are part of one hoop), while tetrameric cross-links must connect four glycan strands (two in one hoop and two in an adjacent hoop). *PG_maker* then determines where to place those Lpp trimers that contain more than one covalent connection to the PG. In effect, such Lpps form an additional source of cross-links since they cause glycan strands on adjacent hoops to be connected due to their shared covalent connections to an intervening Lpp. As noted earlier, the code allows Lpps to be connected to two or three free peptide sidechains, all of which we assume must be on separate glycan strands. Next, the code determines where to add the large numbers of regular dimeric peptide cross-links that connect glycan strands in adjacent hoops; as noted above, the code allows three different types of dimeric cross-link to be added (tetra-DD-tetra, tetra-DD-tri, and tetra-LD-tri), and while they appear identical in the highly idealized models produced by *PG_maker* in this manuscript, they are labeled differently to facilitate their different chemical compositions being respected in future dynamics simulations.

The process of determining where to place all peptide cross-links is carried out stochastically until the desired numbers have been added or until no further cross-links can be added due to geometric constraints. The code selects candidate MurNAc residues for cross-linking randomly, and then seeks to determine whether there are cross-link partners on adjacent glycan strands for which the peptide side-chains are suitably oriented (i.e. pointing toward each other) and for which the glycans are within a specified cutoff distance (set to 35 Å in the models reported here). If all geometric criteria are fulfilled, the cross-link is considered successfully placed and, if necessary, the sidechain orientations of all other MurNAc residues in the involved glycan strands are set. Having attempted to match all of the user’s desired numbers of the various cross-link types, *PG_maker* then provides an option for placing small numbers of additional dimeric cross-links so as to ensure that every glycan strand in the model contains at least one cross-link: without invoking this option it is possible that glycan strands might, by sheer luck, have been left completely neglected during the stochastic process of cross-link incorporation. Finally, since the latter option only seeks to ensure that glycan strands contain at least one cross-link, it is still possible for a strand to contain long, uninterrupted stretches of uncross-linked MurNAc residues. The code therefore also provides an option that seeks to add further cross-links to prevent this from happening. In the models presented here, this option was invoked to ensure that strands have no more than 4 contiguous uncross-linked MurNAc residues.

Once all decisions have been made about the placement of the various types of peptide cross-link, the actual 3D model-building of the peptides can begin in earnest. First, the coordinates of sidechains that are uncross-linked but that are either free or covalently connected to a singly-connected Lpp are built; to ensure that free sidechains on adjacent glycan hoops do not clash sterically, the coordinates of the sidechain CG beads are displaced slightly in a direction pointing toward the center of the cell. Next, the coordinates of all CG beads involved in cross-links are determined. Finally, the coordinates of all Lpps that contain more than one covalent connection to the PG are constructed. Reflecting its rod-like coiled-coil shape (see Figure S5) Lpp itself is modeled as a linear sequence of nine CG beads, each with a radius of 10 Å and separated from the next CG bead by ∼10.6 Å; all Lpps are assumed at this stage to be oriented perpendicular to the plane of the PG and directed so that their N-termini (which in reality are tripalmitoylated) sit in the inner-leaflet of the outer membrane.

As noted above, *PG_maker*’s decisions about the order in which glycans are to be considered possible sites for adding cross-links are made stochastically. The geometric restrictions associated with certain cross-links can mean that their output numbers can be lower than requested by the user and this is particularly true of the tetrameric peptide cross-link (see Results). On the other hand, the code’s options for placing additional dimeric cross-links so as to ensure that no glycan strand is left uncross-linked can cause the output numbers to sometimes be greater than requested by the user. The under- or over-shooting of the desired abundances are generally small – typically less than 5% – so the models produced by a single run of the *PG_maker* code usually provide an accurate realization of the PG composition requested by the user. Stricter adherence to the desired abundances can be enforced simply by running the *PG_maker* code iteratively, continually adjusting the input abundances until the desired output is achieved. For the models reported here, this process was conducted manually, but in the future, it should be possible to implement an automated procedure.

### Quantifying pore sizes in 3D PG models

To identify pores within the PG and to quantify their areas, a number of different images of each final PG model were rendered in VMD (61) with the CG beads of the PG colored in white and all Lpp beads hidden. The resolution of these images was set to 1008 × 1008 pixels. For the quantification, 4 replicate whole-cell models of the PG were generated using the same input parameters but with different random seeds used to start the model-building process. Images were rendered of the cylindrical and polar regions of each PG model and combined in suitably weighted form to yield statistics for the whole cell. Each image was processed in ImageJ/Fiji (44,45) to convert them to 8-bit grey scale, and the pores were quantified with the “auto threshold” tool using default parameters. The commands for processing the images within ImageJ were as follows:

~~~
arg = getArgument();
File.openSequence(arg);
run(“Set Measurements…”, “area perimeter fit stack display redirect=None decimal=3”); run(“8-bit”);
run(“Auto Threshold”, “method=Default stack”);
run(“Analyze Particles…”, “display exclude summarize stack”); saveAs(“Results”, “Results.csv”);
~~~

The output from the above commands is an ordered list of pores that have been identified by the segmentation algorithm. The pores shown in Figures 5a and 5b are colored and labeled using ImageJ’s graphical user interface.

### Computing free energy profiles for proteins crossing the periplasm

To determine the potential thermodynamic effects exerted by the PG and its attached Lpps on the ease with which freely diffusing proteins might cross the periplasm we conducted calculations as follows. First, five representative periplasmic proteins from *E. coli* were selected for study, all of which are monomeric, and for all of which have atomic models are available at the AlphaFold2 (62) database hosted by the EMBL-EBI (https://alphafold.ebi.ac.uk; (63)). Each structure was downloaded and converted into a simplified coarse-grained (CG) representation using the K-means utility `qpdb` that is provided as part of the Situs package (64); all CG models were constructed at a resolution of 10 residues per bead. To identify steric clashes between the protein and the whole-cell PG model, we needed to first assign reasonable radii to each of the CG beads in the combined model. For the proteins, the radius of each bead was assigned automatically by `qpdb` based on the number of atoms assigned by the K-means algorithm to the bead. For the beads of the PG, radii were based on exploratory molecular dynamics simulations that we have performed for a variety of peptidoglycan fragments. For the beads of Lpp, we assigned a constant radius of 10 Å to match the approximate radius of the atomic structure (49) (RCSB ID: 1EQ7; Figure S5).

The free energy profile for translating each protein across the periplasm was then calculated using additional computer code as follows. The protein was randomly rotated, and its center placed at a randomly selected position on the inner membrane. The protein was then moved in 1 Å intervals in an outward radial direction until it reached the outer membrane; we assume that this involves a movement of 250 Å. At each radial distance, the protein was checked for steric clashes with the PG, the Lpps, and the two membranes. Any orientations that resulted in steric clashes were rejected and a tally of the number of accepted orientations at each radial distance was recorded. This process was repeated 10 million times for each protein, with each new attempt involving a new random rotation of the protein and a newly selected starting position within the inner membrane. The final free energy as a function of radial distance was calculated using ΔG = - k_B_T ln ((N_acc_ + 1) / 10^7^), where k_B_ is Boltzmann’s constant, T is the temperature (set here to 298 K), N_acc_ is the number of random orientations of the protein that were accepted at that radial distance, and 1 is added to the counts to ensure that the computed ΔG is capped at a finite value for those radial distances at which no orientations were accepted (see Results). Separate free energy profiles were computed for the polar and cylindrical regions of the cell but in practice we saw very little difference between them.

### Computing hardware

All simulations and model generation were carried out on compute nodes equipped with Xeon Gold 6130@2.1GHz CPUs and 128 GB RAM. All machines were housed and maintained as a part of the University of Iowa’s Argon high performance computing cluster.

### Figure generation

Bar charts, histograms, and scatterplots were made in the statistical programming language R (R Core Team, 2023) with the following additional packages: ggplot2 (Wickham, 2022), gridExtra (Auguie, 2017), and cowplot (Wilke, 2020). Images of the PG and proteins were made with VMD (61).

## Acknowledgments

This research was supported by a grant from the National Institutes of Health (R35 GM122466) to AHE and supported in part through computational resources provided by The University of Iowa. Both authors thank Professor David S. Weiss (University of Iowa) for very valuable discussions.

## Author Contributions (CRediT)

Zachary J. Wehrspan: Methodology, Software, Formal analysis, Investigation, Writing – original draft preparation, Visualization.

Adrian H. Elcock: Conceptualization, Methodology, Software, Formal analysis, Investigation, Writing – review and editing, Supervision, Project Administration, Funding acquisition.

## Data availability

All computer code necessary to run the simulations described here will be made available to reviewers at the time of manuscript review. Upon acceptance of the manuscript for publication, the computer code will be available to the community at the following GitHub repository (https://github.com/Elcock-Lab/peptidoglycan).

