## Supporting Information for "Multi-scale Modeling and Experimental Data Enable Structural Models of the *Escherichia Coli* Peptidoglycan to be Constructed on the Whole-Cell Scale"

### Supporting Figure Legends:

**Figure S1** Changing the assumed standard deviation of the Gaussian function describing the long glycan strand population causes little to change to its optimized mean position. The polar strand length distributions are colored in blue, and the cylindrical distributions are colored in blue. For both, three optimized distributions are plotted, with standard deviations (Std) of 15 DS units (light blue), 30 DS units (medium blue), and 60 DS units (dark blue).

**Figure S2** Integrated intensities of bands on the immunoblot shown in the highest DSS concentration lanes in Figure 6 of Cowles et al. Quantification was performed as described in the main text using ImageJ/Fiji. (a) wildtype Lpp. (b) $\Delta K_{58}$ mutant.

**Figure S3** Variation in the relative populations of DSS cross-linked oligomers derived from the integrated intensities shown in Figure S2. Each panel illustrates a different possible partitioning of the immunoblot intensities for determining the relative DSS cross-linked monomer, dimer, and trimer Lpp populations. The first 6 panels show results for the wild-type Lpp data shown in Figure S2a; the final panel shows results for the $\Delta K_{58}$ mutant shown in Figure S2b. The left-hand side of each panel shows the lines used to partition the plot by oligomer size, with the relative areas within each partition also displayed; the right-hand side of each panel shows the resulting best agreement obtained from stochastic simulations for each of the relative DSS cross-linked populations (using the same coloring scheme used in Figure 3).

**Figure S4** A model in which all Lpps are located in the periplasm can capture experiment as well as other models if ~17% of the periplasmic Lpp trimers are assumed to be unconnected to the PG. Using the same display scheme used in Figures 4 and S3, results are shown for three different possible scenarios in which 100% of the Lpps are assumed to be resident in the periplasm. The ratios on the x-axis indicate, respectively, the relative numbers of Lpp trimers that are unconnected, singly-connected, doubly-connected, and triply-connected to the PG. For example, the scenario shown at the far-right (given the ratio 2:8:2:0) contains 17% unconnected, 67% singly-connected, and 17% doubly-connected Lpps.

**Figure S5** Image of the complete atomic model of the Lpp trimer homology-modeled using in-house code and based on the crystal structure (RCSB ID: 1EQ7). The protein is oriented such that the N-termini of all three chains are at the top of the image. The protein is shown in licorice representation; the lysine sidechains are shown in the following CPK representations: K6 (blue), K20 (cyan), K38 (green), K55 (yellow), K58 (red).

**Figure S6** Iterative optimization of the error in the output glycan strand length distributions obtained from *PG_maker*. For reasons outlined in the main text, the length distributions output by the code can deviate from the length distributions that are input to the code. A simple iterative scheme can be used to separately adjust the polar (red) and cylindrical (blue) length distributions until the output distributions match the desired distributions.

**Figure S7** Same as Figure 1 of the main text but comparing the glycan strand length distributions before (dashed lines) and after (solid lines) the optimization process used to ensure that the final distributions output by *PG_maker* match the desired distributions.

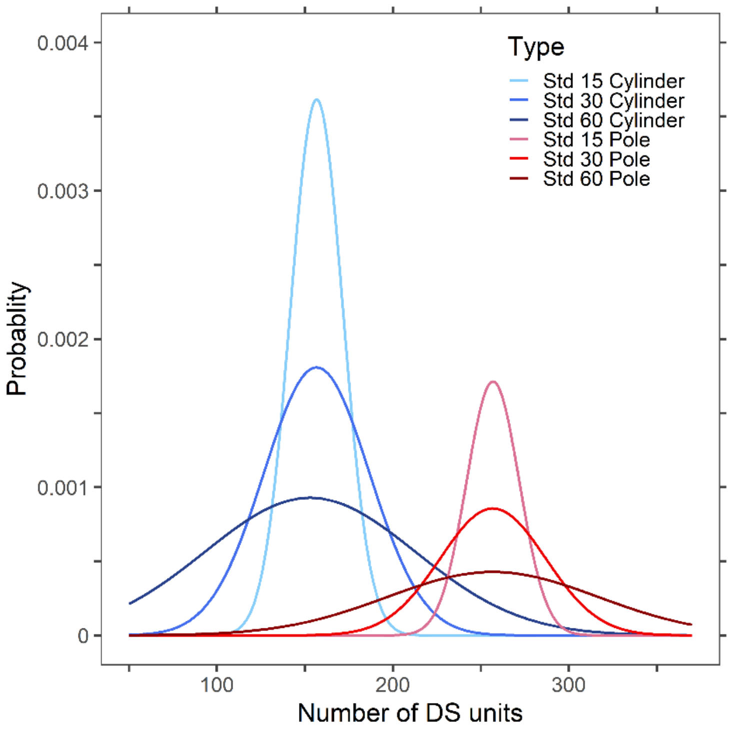
Type
Std 15 Cylinder
Std 30 Cylinder
Std 60 Cylinder
Std 15 Pole
Std 30 Pole
Std 60 Pole
Probablity
0.004
0.003
0.002
0.001
0
100
200
300
Number of DS units

**Figure S1**

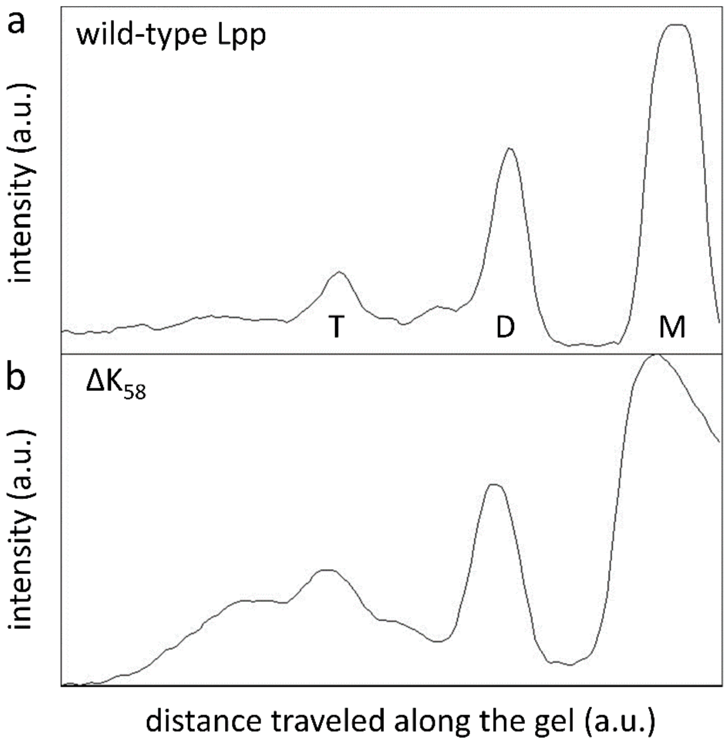
a
wild-type Lpp
intensity (a.u.)
T
D
M
b
ΔK58
intensity (a.u.)
distance traveled along the gel (a.u.)

**Figure S2**

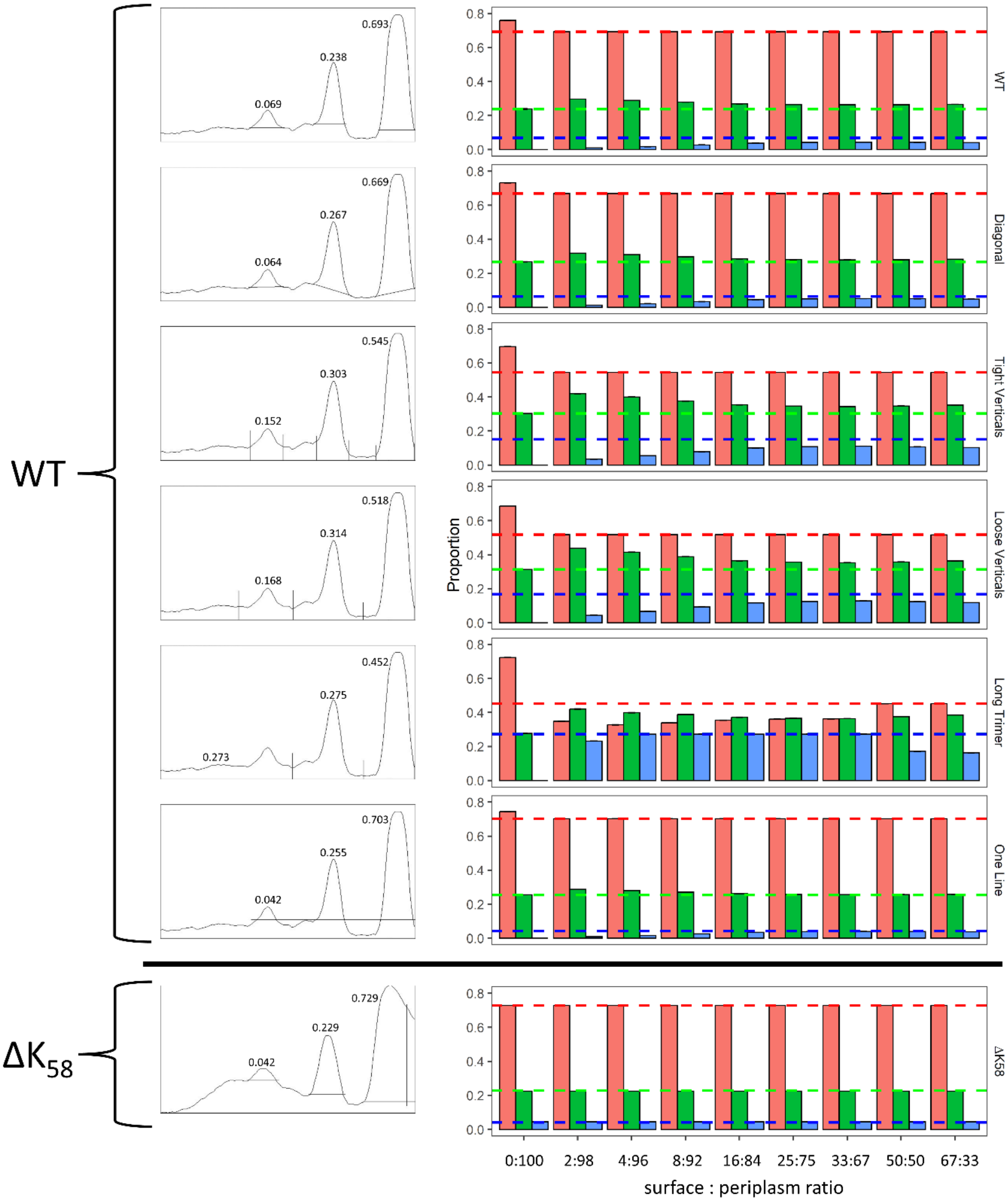
WT
ΔK58
0.693
0.238
0.069
0.669
0.267
0.064
0.545
0.303
0.152
0.518
0.314
0.168
0.452
0.275
0.273
0.703
0.255
0.042
0.729
0.229
0.042
Proportion
WT
Diagonal
Tight Verticals
Loose Verticals
Long Trimer
One Line
ΔK58
0:100
2:98
4:96
8:92
16:84
25:75
33:67
50:50
67:33
surface : periplasm ratio

**Figure S3**

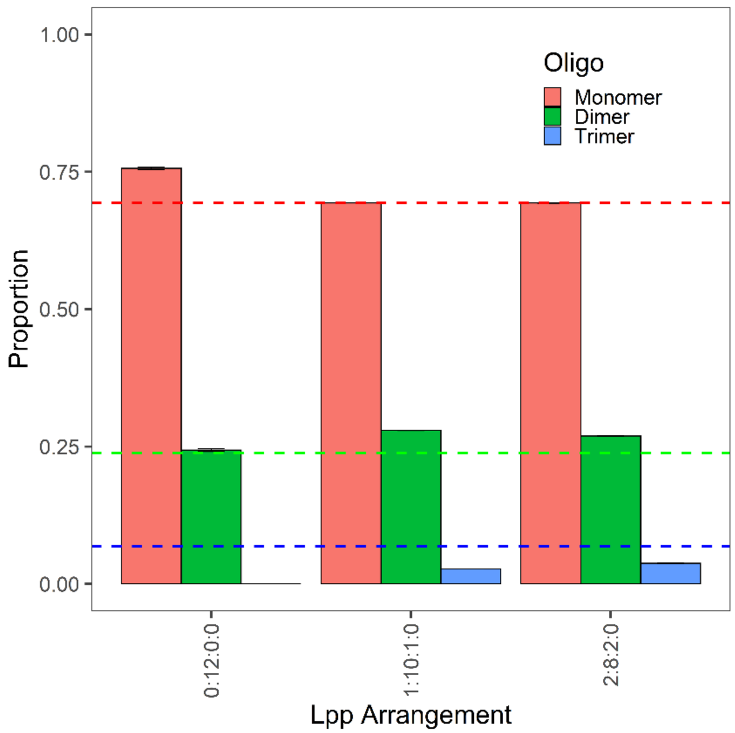

Oligo
Monomer
Dimer
Trimer
Proportion
1.00
0.75
0.50
0.25
0.00
0:12:0:0
1:10:1:0
2:8:2:0
Lpp Arrangement


**Figure S4**

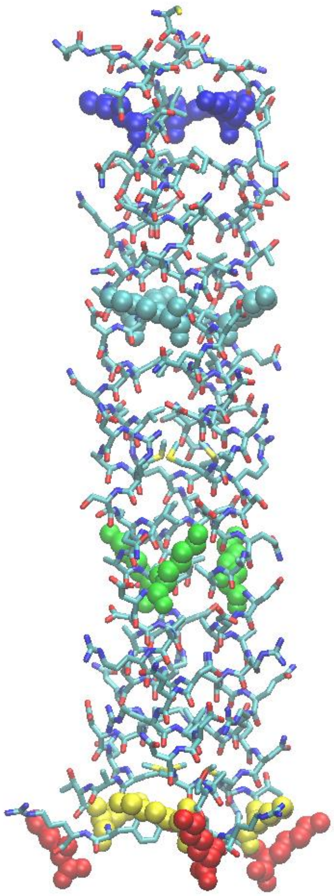

**Figure S5**

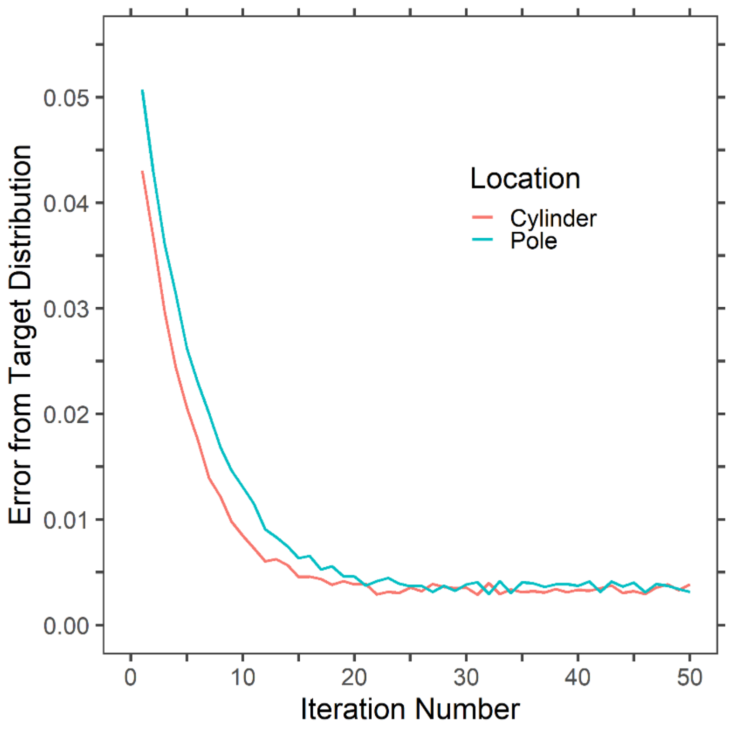

Error from Target Distribution
0.05
0.04
0.03
0.02
0.01
0.00
0
10
20
30
40
50
Iteration Number
Location
Cylinder
Pole


**Figure S6**

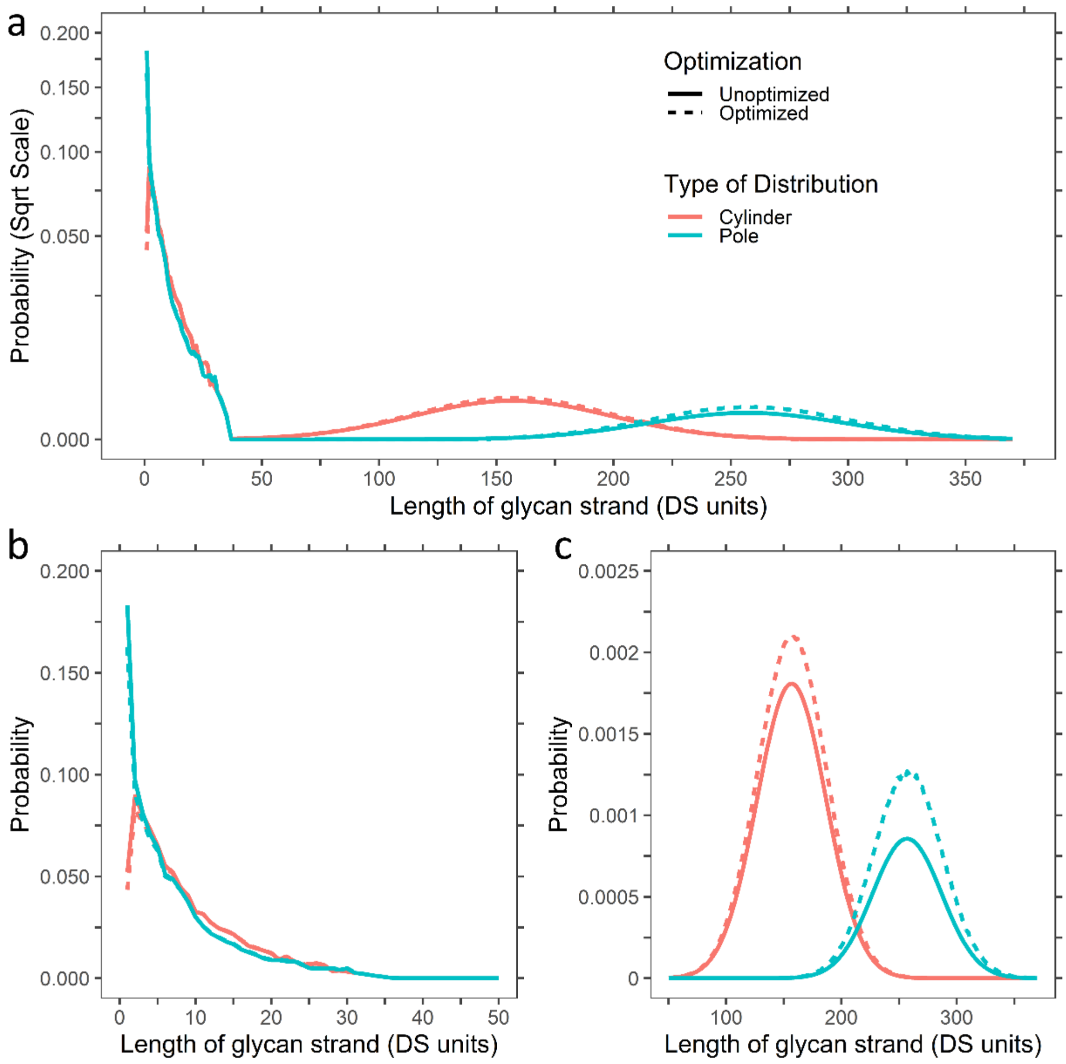
a
Probability (Sqrt Scale)
0.200
0.150
0.100
0.050
0.000
Optimization
Unoptimized
Optimized
Type of Distribution
Cylinder
Pole
0
50
100
150
200
250
300
350
Length of glycan strand (DS units)
b
Probability
0.200
0.150
0.100
0.050
0.000
0
10
20
30
40
50
Length of glycan strand (DS units)
c
Probability
0.0025
0.002
0.0015
0.001
0.0005
0
100
200
300
Length of glycan strand (DS units)

**Figure S7**
